# ANANSE: An enhancer network-based computational approach for predicting key transcription factors in cell fate determination

**DOI:** 10.1101/2020.06.05.135798

**Authors:** Quan Xu, Georgios Georgiou, Siebren Frölich, Maarten van der Sande, Gert Jan C. Veenstra, Huiqing Zhou, Simon J. van Heeringen

## Abstract

Proper cell fate determination is largely orchestrated by complex gene regulatory networks centered around transcription factors. However, experimental elucidation of key transcription factors that drive cellular identity is currently often intractable. Here, we present ANANSE (<u>AN</u>alysis <u>A</u>lgorithm for <u>N</u>etworks <u>S</u>pecified by <u>E</u>nhancers), a network-based method that exploits enhancer-encoded regulatory information to identify the key transcription factors in cell fate determination. As cell type-specific transcription factors predominantly bind to enhancers, we use regulatory networks based on enhancer properties to prioritize transcription factors. First, we predict genome-wide binding profiles of transcription factors in various cell types using enhancer activity and transcription factor binding motifs. Subsequently, applying these inferred binding profiles, we construct cell type-specific gene regulatory networks, and then predict key transcription factors controlling cell fate transitions using differential networks between cell types. This method outperforms existing approaches in correctly predicting major transcription factors previously identified to be sufficient for trans-differentiation. Finally, we apply ANANSE to define an atlas of key transcription factors in 18 normal human tissues. In conclusion, we present a ready-to-implement computational tool for efficient prediction of transcription factors in cell fate determination and to study transcription factor-mediated regulatory mechanisms. ANANSE is freely available at https://github.com/vanheeringen-lab/ANANSE.

## Introduction

Every multicellular organism develops from a single cell. During this process, cells undergo division and differentiation, eventually forming a diversity of cell types that are organized into organs and tissues. How one cell develops into different cell types, a process known as cell fate determination, is critical during development. It has been shown that transcription factors (TFs) play key roles in cell fate determination (Davis et al., 1987; Jopling et al., 2011; Pang et al., 2011; Stadhouders et al., 2019; Takahashi et al., 2007; Vierbuchen et al., 2010). TFs bind to specific cis-regulatory sequences in the genome, including enhancers and promoters, and regulate expression of their target genes (Lambert et al., 2018; Vaquerizas et al., 2009). The interactions between TFs and their downstream target genes form gene regulatory networks (GRNs), controlling a dynamic cascade of cellular information processing which shapes cell fate and identity (Davidson, 2010; Tegner and Bjorkegren, 2007). Cell fate determination is orchestrated by a series of TF regulatory events, largely by complex GRNs (Wilkinson et al., 2017). The key role of TFs and GRNs in cell fate determination is further corroborated by examples of cell fate conversions, often referred to as cellular reprogramming (Iwafuchi-Doi and Zaret, 2016; Peñalosa-Ruiz et al., 2019). Cellular reprogramming includes generating induced pluripotent stem cells (iPSCs) from somatic cells, and trans-differentiation that converts one mature somatic cell type to another without undergoing an intermediate pluripotent state (Davis et al., 1987; Jopling et al., 2011; Pang et al., 2011; Stadhouders et al., 2019; Takahashi et al., 2007; Vierbuchen et al., 2010). These reprogramming processes are initiated by enforced expression of combinations of key TFs, which is believed to alter the GRNs at the level of gene expression and the epigenetic landscape (Buschbeck and Hake, 2017; Qu et al., 2018; Reik et al., 2001).

In the past, identification of key TFs driving cellular differentiation or reprogramming was often performed by experimental screening or testing candidate genes, which is labor-intensive and inefficient. Therefore, there is a need for better predictions of key TFs in cell fate determination, which can help to understand developmental processes and serve to instruct experimental cellular reprogramming approaches. Different computational methods for predicting key transcription factors or master regulators in the context of cellular transitions have been reported. Some are based on gene expression differences between cell types (D’Alessio et al., 2015; Heinaniemi et al., 2013; Lang et al., 2014; Roost et al., 2015). Other methods use GRNs in combination with expression differences to identify candidate key TFs (Alvarez et al., 2016; Cahan et al., 2014; Hartmann et al., 2018; Morris et al., 2014; Rackham et al., 2016). However, these GRNs are usually inferred based on a measure of co-expression (Huynh-Thu et al., 2010; Margolin et al., 2006), which requires many different samples and which cannot easily distinguish directionality.

The use of (predicted) transcription factor binding sites allows for directionality and has been shown to improve GRN inference (Glass et al., 2013; Janky et al., 2014; Marbach et al., 2012). Mogrify is an example of a method that uses not only gene expression but also GRNs constructed based on TF binding motifs in promoters to predict TFs that are capable of inducing conversions between cell types (Rackham et al., 2016). However, most GRN-based approaches that incorporate TF motifs only include promoters or promoter-proximal regulatory elements. It has been well established that TFs that control tissue-and cell type-specific gene expression in cell fate determination and development often bind to enhancers (Andersson et al., 2014; Bulger and Groudine, 2011; Qu et al., 2018; Spitz and Furlong, 2012). Binding of tissue-and cell type-specific TFs largely to enhancers is also confirmed by a large number of genome-wide chromatin immunoprecipitation followed by sequencing analyses (ChIP-seq) (Davis et al., 2018; Valouev et al., 2008), e.g. TP63 in keratinocytes and ZIC2 in embryonic stem cells (Luo et al., 2015; Qu et al., 2018). Furthermore, analysis of enhancers and enhancer clusters allows for identification of master regulators, corroborating their relevance in cell type-specific gene regulation (Huang et al., 2017; Whyte et al., 2013). Large compendia of transcription factor binding profiles and enhancer-associated histone modifications can also be used to prioritize transcriptional regulators (Qin et al., 2020; Wang et al., 2018). Therefore, we reasoned that a computational method that uses enhancer properties to infer enhancer-based GRNs would improve the prediction of directed regulatory interactions at a genome-wide scale. Furthermore, most current computational tools require comprehensive training or background data, such as cell/tissue expression data or pre-constructed networks. This means that they cannot be applied in new biological contexts or in non-model species that are less well-studied. Finally, these datasets and the computational algorithms are not always publicly accessible, which prevents the general usage of these methods in studying transcriptional regulation or designing new trans-differentiation strategies.

Here, we established an enhancer GRN-based method, ANalysis Algorithm for Networks Specified by Enhancers (ANANSE), that infers genome-wide regulatory programs and identifies key TFs for cell fate determination. We predicted cell type-specific TF binding profiles with a model that incorporates activities and sequence features of enhancers. Second, combining TF binding profiles and gene expression data, we built cell type-specific enhancer GRNs in each cell type or tissue. We used reference GRNs, constructed from known TF-target gene interactions and experimental data of TF perturbations, to evaluate the quality of the inferred GRNs. Third, we predicted the key TFs underlying cell fate conversions based on a differential network analysis. Compared with other reported prediction algorithms, ANANSE recovers the largest fraction of TFs that were validated by experimental trans-differentiation approaches. The results demonstrate that ANANSE can prioritize TFs that drive cellular fate changes. Finally, to demonstrate the wide utility of ANANSE, we applied it to 18 human tissues and generated an atlas of key TFs underlying human tissue identity.

## Materials and Methods

### Analysis of the genomic distribution of TF binding sites

For every transcription factor, we combined all the peaks in the ReMap database (Chèneby et al., 2017) by taking the peaks in all cell types and tissues for this specific TF. TFs that had less than 600 peaks were removed. This resulted in a data set of ChIP-seq peaks from 296 unique TFs. The percentage of peaks in each genomic location was calculated using the ChIPseeker R package (version 1.20.0) (Yu et al., 2015). The fgsea R package (version 1.10.1) was used to do the gene set enrichment analysis (GSEA) (Sergushichev, 2016).

We used the classification of the Human Protein Atlas (Uhlén et al., 2015) to determine tissue-specific TFs. This classification is divided in several groups based on gene expression patterns using RNA-seq from human tissues. We took the union of tissue enriched genes (at least a 5-fold higher FPKM level in one tissue compared to all other tissues), group enriched genes (5-fold higher average FPKM value in a group of 2-7 tissues compared to all other tissues) and tissue enhanced genes (at least a 5-fold higher FPKM level in one tissue compared to the average value of all 32 tissues).

### Datasets

All H3K27ac ChIP-seq, ATAC-seq, and RNA-seq data used in this study was obtained from GEO or ENCODE (Bernstein et al., 2010; Buenrostro et al., 2018; Cho et al., 2018; Encode Project Consortium, 2012; Granja et al., 2021; Grubert et al., 2020; Johnston et al., 2019; Li et al., 2019b; Liu et al., 2017; Liu et al., 2020; Martone et al., 2020; Morris et al., 2019; Novakovic et al., 2016; Oomen et al., 2019; Runge et al., 2018; Segura-Bayona et al., 2020; Soares et al., 2019; Song et al., 2019; Tchieu et al., 2019). For all data sets with ENCODE identifiers we downloaded the BAM files (ATAC-seq; H3K27ac ChIP-seq) or the FASTQ files (RNA-seq) from the ENCODE portal (Sloan et al., 2016) (https://www.encodeproject.org/). For data sets with a GSM accession, FASTQ files were downloaded and further processed using seq2science (version v0.4.3) (van der Sande et al., 2020), see paragraph below. All data sets and accession numbers are summarized in Supplementary Table S1.

### Data and code availability

ANANSE source code is available from https://github.com/vanheeringen-lab/ANANSE. Jupyter notebooks for supporting analyses are provided at https://github.com/vanheeringen-lab/ANANSE-manuscript). GRNs inferred with ANANSE are available from Zenodo (tissue-specific GRNs https://doi.org/10.5281/zenodo.4814016; cell type-specific GRNs https://doi.org/10.5281/zenodo.4809062). Tissue-specific GRNs inferred with GRNBoost2 are available from Zenodo (https://doi.org/10.5281/zenodo.4814015).

### ChIP-seq, ATAC-seq and RNA-seq analyses

Analysis of publicly available ChIP-seq, ATAC-seq and RNA-seq analysis was performed with seq2science (version v0.4.3) (van der Sande et al., 2020). Genome assembly hg38 was downloaded from UCSC with genomepy 0.9.1 (van Heeringen, 2017). The reads of the ChIP-seq experiments were mapped to the human genome (hg38) using STAR (version 2.5.3a) with default settings (Dobin et al., 2013). Duplicate reads were marked and removed using Picard (Picard2019toolkit, 2019). Peaks were called on the ChIP-seq data with only the uniquely mapped reads using MACS2 (version 2.7) relative to the Input track using the standard settings and a q-value of 0.01 (Zhang et al., 2008). The measurement of consistent peaks between replicates was identified by IDR (version 2.0.3) (Li et al., 2011).

ATAC-seq reads were trimmed with fastp v0.20.1 (Chen et al., 2018) and aligned with bwa-mem v0.7.17 (Li, 2013) to the hg38 genome. Mapped reads were removed if they did not have a minimum mapping quality of 30, were a (secondary) multimapper or aligned inside the ENCODE blacklist (Amemiya et al., 2019). Reads were shifted for tn5 bias. Duplicate reads were removed with picard MarkDuplicates v2.23.8 (Picard2019toolkit, 2019). Peaks were called with macs2 v2.2.7 (Zhang et al., 2008) with options ‘--shift -100 --extsize 200 --nomodel --keep-dup 1 --buffer-size 10000’ in BAM mode.

Quantification of expression levels was performed on RNA-seq data, using salmon (version 0.43.0) (Patro et al., 2017) with default settings and Ensembl transcript sequences (version GRCh38 release-103) (Cunningham et al., 2018). Salmon’s transcript-level quantifications results were imported and aggregated to gene level counts by the tximport R package (version 1.12.3) (Soneson et al., 2015). The expression level (transcript-per-million, TPM) of each cell type and the differential expression fold change between two cell types were calculated using the DESeq2 R package (version 1.24.0) (Love et al., 2014). The expression TPM data used to predict key TFs for trans-differentiation is shown in Supplementary Table S3 and differential gene expression data is shown in Supplementary Table S4.

### Defining putative enhancer regions

To generate a collection of putative enhancer regions, we collected all transcription factor ChIP-seq peaks from ReMap 2018 (http://remap.univ-amu.fr/storage/remap2018/hg38/MACS/remap2018_all_macs2_hg38_v1_2.bed.gz) (Chèneby et al., 2017). We took the summit of all peaks and extended these 25 bp up-and downstream. Based on this file, we generated a coverage bedGraph using bedtools genomecov (Quinlan and Hall, 2010). We performed peak calling on this bedGraph file using bdgpeakcall from MACS2 (version v2.7.1) (Zhang et al., 2008), with the following settings: l=50 and g=10. We performed the peak calling twice, setting c to 4 and 30, respectively. All peaks from c=30 were combined with all peaks of c=4 that did not overlap with the peaks of c=30. We then removed all regions on chrM and extended the summit of the peaks 100 bp up-and downstream to generate a final collection of 1,268,775 putative enhancers of 200 bp. This collection of enhancers is available at Zenodo with doi 10.5281/zenodo.4066423.

The coverage_table script from GimmeMotifs 0.15.3 (Bruse and Heeringen, 2018; van Heeringen and Veenstra, 2010) was used to determine the ATAC-seq and H3K27ac intensity, as expressed by the number of reads, in all enhancer peaks (2,000 bp centered at the enhancer summit for H3K27ac; 200bp for ATAC-seq). All counts were quantile normalized using qnorm v0.4.0 (van der Sande and van Heeringen, 2021).

### Prediction of transcription factor binding

To train the ANANSE models we used ChIP-seq peaks for 237 TFs from REMAP in 6 cell types: hESC, Hep-G2, HeLa-S3, K562, MCF-7 and GM12878. ATAC-seq and H3K27ac ChIP-seq data for these cell types was downloaded from public repositories, see Supplementary Table S1. For both assays, the number of reads was determined in regions of 200 bp (ATAC-seq) or 2kb (H3K27ac) centered at the enhancer summit. Read counts were log-transformed and quantile normalized. To test the prediction performance of the ANANSE model a cross-validation procedure was used. For each TF, models were trained on binding in all enhancers, except those on chromosomes chr1, chr8 and chr23 (held-out chromosomes). The evaluation was only performed on those TFs for which peaks in multiple cell types were available. Each cell type was left out (held-out cell types) and the classifier was trained on data of the other cell type(s). In this manner, performance metrics (ROC AUC and PR AUC) were calculated based on enhancers located on held-out chromosomes in held-out cell types.

Binding was predicted using four type(s) of input features: TF motif scores, ATAC-seq signal in enhancers, H3K27ac ChIP-seq signal in enhancers and (optionally) the average ChIP-seq signal of REMAP peaks in enhancers. ANANSE uses a standard logistic regression model as implemented in scikit-learn (Pedregosa et al., 2011). Equation 1 shows an example of a model, using all four types of input.

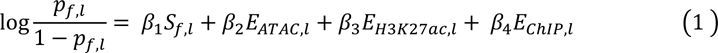

Where *p*_*f*,*l*_ is the probability of a transcription factor *f* binding to enhancer *l*. *S*_*f*,*l*_ is the highest motif z-score of all motifs associated with transcription factor *f* in enhancer *l* and *E*_*ATAC*,*l*_, *E*_*H*3*K*27*ac*,*l*_ and *E*_*C*ℎ*IP*,*l*_ represent the enhancer intensity of enhancer *l*, based on scaled and normalized ATAC-seq signal, scaled and normalized H3K27ac ChIP-seq signal and average REMAP ChIP-seq signal, respectively.

ANANSE incorporates a flexible selection of models, the choice of which depends on the type of input that is available. The minimal input consists of the motif score and either ATAC-seq or H3K27ac ChIP-seq signal in enhancers.

The non-redundant database of 1,796 motifs was created by clustering all vertebrate motifs from the CIS-BP database using GimmeMotifs (van Heeringen and Veenstra, 2010; Weirauch et al., 2014) as described in (Bruse and Heeringen, 2018). The GimmeMotifs package (version 0.15.3) (Bruse and Heeringen, 2018; van Heeringen and Veenstra, 2010) was used to scan for motifs in enhancer regions. The GC normalization setting in GimmeMotifs package was used to normalize the GC% bias in different enhancers. To correct for the bias of motif length, z-score normalization was performed on the motif scores. Normalization was done per motif, based on motif matches in random genomic regions using the same motif scan settings. The highest z-score was chosen if a TF had more than one motif.

### Gene regulatory network inference

The weighted sum of the TF binding probability, predicted on the basis of the enhancer intensity and the motif score, within 100kb around TSS is defined as the TF-gene binding score (Eq. 2). The distance weight is based on a linear genomic distance between the enhancer and the TSS of a gene according to equation 3.

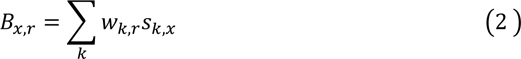

where *B*_*x*,*r*_ is the binding score between TF *x* and target gene *r*, *w*_*k*_ is the weighted distance between an enhancer and the target gene and where *s*_*k*_ is predicted binding intensity at genomic position *k* of TF *x*.

The distance weight calculation was similar to the method previously described in (Wang et al., 2016), except that only the signal in predetermined enhancers is used and the weight of enhancers within 5kb of the TSS is set to 1.

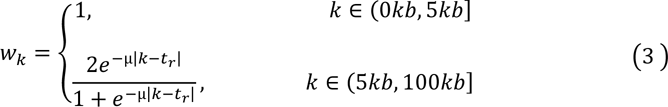

where *t*_*r*_ is the genomic position of the TSS of gene *r* and the parameter µ, which determines the decay rate as a function of distance from the TSS, is set such that an enhancer 10 kb from the TSS contributes one-half of an enhancer within 5kb from TSS.

We determined a measure of genome-wide TF activity, *A*_*x*_, based upon the motif activity. The motif activity for all TF motifs was calculated based on ridge regression as implemented in scikit-learn (Pedregosa et al., 2011) using GimmeMotifs 0.15.3 (Bruse and Heeringen, 2018). Here, the motifs scores were used as input to predict either ATAC-seq and/or H3K27ac ChIP-seq signal. The TF activity is the maximum activity of the motifs associated with a TF, where the motif activity is defined as the mean of the ATAC-seq motif coefficients and the H3K27ac ChIP-seq coefficients

The expression level of the TF *E*_*x*_ and the target gene *E*_*r*_, expressed as transcripts per million (TPM), and the TF activity *A*_*x*_ and TF-gene binding score *B*_*x*,*r*_ were ranked and scaled, from 0 to 1, where 0 represents the lowest value and 1 represents the highest value. For ranking the the TF expression, only the expression levels of TFs where used. The interaction score was calculated (Eq. 4) by mean averaging the individual ranked scores (mean rank aggregation).

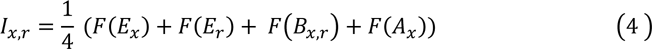

Where *I*_*x*,*r*_ is the interaction score between TF *x* and target gene *r* and *F*(*X*)represents the rank aggregated and scaled score. Ideally, the contributions of these individual scores would be determined by a supervised method, such as a linear regression, however, due to the lack of a high-quality gold standard reference data set we chose to combine the scores through mean averaging.

### Gene regulatory network evaluation

We obtained GRNs from different sources. GRNBoost2, as implemented in arboreto 0.1.5, was used with default settings to infer networks from GTEx data. The GTEx expression data (GTEx_Analysis_2017-06-05_v8_RNASeQCv1.1.9_gene_tpm.gct.gz) was downloaded from the GTEx portal (https://www.gtexportal.org/home/datasets). Tissue-specific PANDA networks were downloaded from https://sites.google.com/a/channing.harvard.edu/kimberlyglass/tools/gtex-networks. Tissue-specific networks inferred from single cell data using SCENIC (Aibar et al., 2017) where downloaded from http://www.grndb.com/ (Fang et al., 2021). GRNs inferred from GTEx data with corto and ARACNE were downloaded from https://giorgilab.org/corto-the-correlation-tool/ (Margolin et al., 2006; Mercatelli et al., 2020)

To evaluate the quality of the predicted GRNs, four different types of reference datasets were used: gene co-expression, Gene Ontology (GO) annotation (The Gene Ontology, 2019), four regulatory interaction databases (DoRothEA (Holland et al., 2020), RegNetwork (Liu et al., 2015), TRRUST (Han et al., 2017) and MSigDB C3 (Liberzon et al., 2011)) and differential expression measurements after TF perturbations. The expression correlation database was downloaded from COXPRESdb (Obayashi et al., 2019), and the original mutual rank correlation score was scaled to 0 to 1 for each TF, with 1 being the highest and 0 the lowest, and all scaled correlation score higher than 0.6 or 0.8 were considered as true interaction pairs. The human GO validation Gene Association File (GAF) (version 2.1) was downloaded from http://geneontology.org. We used all TF-gene pairs that were annotated with at least one common GO term as true positives. The TF perturbation data set was obtained by downloading the “TF_Perturbations_Followed_by_Expression” data set from Enrichr (Chen et al., 2013; Kuleshov et al., 2016). For the random network we used the same network interaction structure, but here we randomized the interaction score (the edge weight) by permutation of the scores. The AUC of ROC and PR for each cell type GRN and corresponding random GRN were calculated.

### Influence score inference

To calculate the influence score for the transition from a source cell type to a target cell type, we used the GRNs for both cell types. In each network, we selected the top 100k interactions based on the rank of its interaction score. We obtained a differential GRN by taking the interactions only located in the target cell type. The difference of the interaction score was used as the edge weight for the differential GRN.

Based upon the differential GRN a local network was built for each TF, up to a maximal number of three edges. Using Equation 5, a target score was calculated for each node in the network, based on 1) its edge distance from the TF of interest, 2) the interaction score and 3) the change in expression between the source cell type and the target cell type.

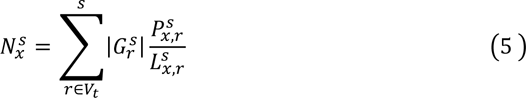

Where *r* ∈ *V*_*t*_ is each gene (*r*) in the set of nodes (*V*_*t*_) that make up the local sub-network of TF *x*. In other words, *V*_*t*_ represents all target genes that are directly or indirectly targeted by TF *x*. To incorporate indirect target genes, only genes up to three steps away are considered. This distance (number of edges or steps) is represented by *L*^*s*^, the level (or the number of steps) that gene *r* is away from TF *x* in the network *s*. Nodes located further from the TF have less effect on the target score. *P*^*s*^ is the interaction score between TF *x* and target gene *r* and *G*_r_ ^*s*^, the expression score, is the log-transformed fold change of the expression of gene *r*.

The target score (*N*^*s*^) for each TF is the sum of the scores from all the nodes in its local network. Nodes present in multiple edges are calculated only for the edge closest to the TF of interest. Self-regulating nodes are not considered. The target score and the *G*^*s*^ of each TF are scaled to 0 to 1, and the mean of them was defined as the influence score of this TF. Subsequently, all TFs are ranked by their influence score.

### Trans-differentiation evaluation

To evaluate the performance the ANANSE influence score calculation we used key TFs from trans-differention experiments. We compared ANANSE results to previously reported methods: Mogrify, LISA, BART, VIPER, CellNet, and the method of D’Alessio et al (Alvarez et al., 2016; Cahan et al., 2014; D’Alessio et al., 2015; Qin et al., 2020; Rackham et al., 2016; Wang et al., 2018). Mogrify and Mogrify full prediction results were downloaded from https://mogrify.net/. For LISA (version 1.2), all differentially expressed genes from fibroblast to each target cell type were used as input (Qin et al., 2020). For BART, we uploaded the top 1000 differentially expressed genes to http://bartweb.org/, using BART 2.0. The DoRothEA network was downloaded from https://github.com/saezlab/dorothea. All differentially expressed genes from fibroblast to each target cell type and networks (ANANSE or DoRothEA) were used as input of VIPER (version 1.24.0). The CellNet predictions were obtained from (Rackham et al., 2016). The prediction results of the method of D’Alessio et al were obtained from the original paper (D’Alessio et al., 2015). Results were compared with the experimentally validated TFs as true positives and all other TFs as false positives.

### Regulatory profile analysis of human tissues

The RNA-seq data of 18 human tissues was downloaded from (https://www.proteinatlas.org/humanproteome/tissue) (Uhlen et al., 2010). The H3K27ac ChIP-seq and ATAC-seq accession numbers are listed in Supplementary Table S1. The gene expression score of each tissue was calculated by taking the log2 TPM fold change between a tissue and the average of all other tissues. The GRN of each tissue was inferred using ANANSE. For prediction for TFs of one tissue, GRN interaction scores of all other tissues were averaged as the source GRN. All correlation analyses were clustered by hierarchical clustering method. The modular visualization of anatograms and tissues was done using the gganatogram package (version 1.1.1)(Maag, 2018).

## Results

### Cell type-specific transcription factors predominantly bind to enhancers

To systematically examine TF binding patterns in the genome in relation to cell type specificity, we downloaded the binding sites of 296 human TFs from the ReMap project, which re-analyzed all publicly available ChIP-seq data in various cell types and tissues (Chèneby et al., 2017). To determine the genomic distribution of these binding sites, we divided the genome into different genomic categories according to human UCSC known gene annotation (Hsu et al., 2006), and assigned binding sites to these categories based on the locations of the binding sites (Figure 1). We grouped these categories into two main classes: 1) a promoter-proximal class, containing promoter (<=2kb), 5’ UTR and 1st exon peaks, and 2) a promoter-distal class, referred to “Enhancers”, containing all exons except the first, the 1st intron, other introns and intergenic categories. The percentage of TF binding sites in each genomic category was calculated, and TFs were ordered according to their percentages in the promoter-proximal class (Figure 1A) (Supplementary Table S2).

**Figure 1.**
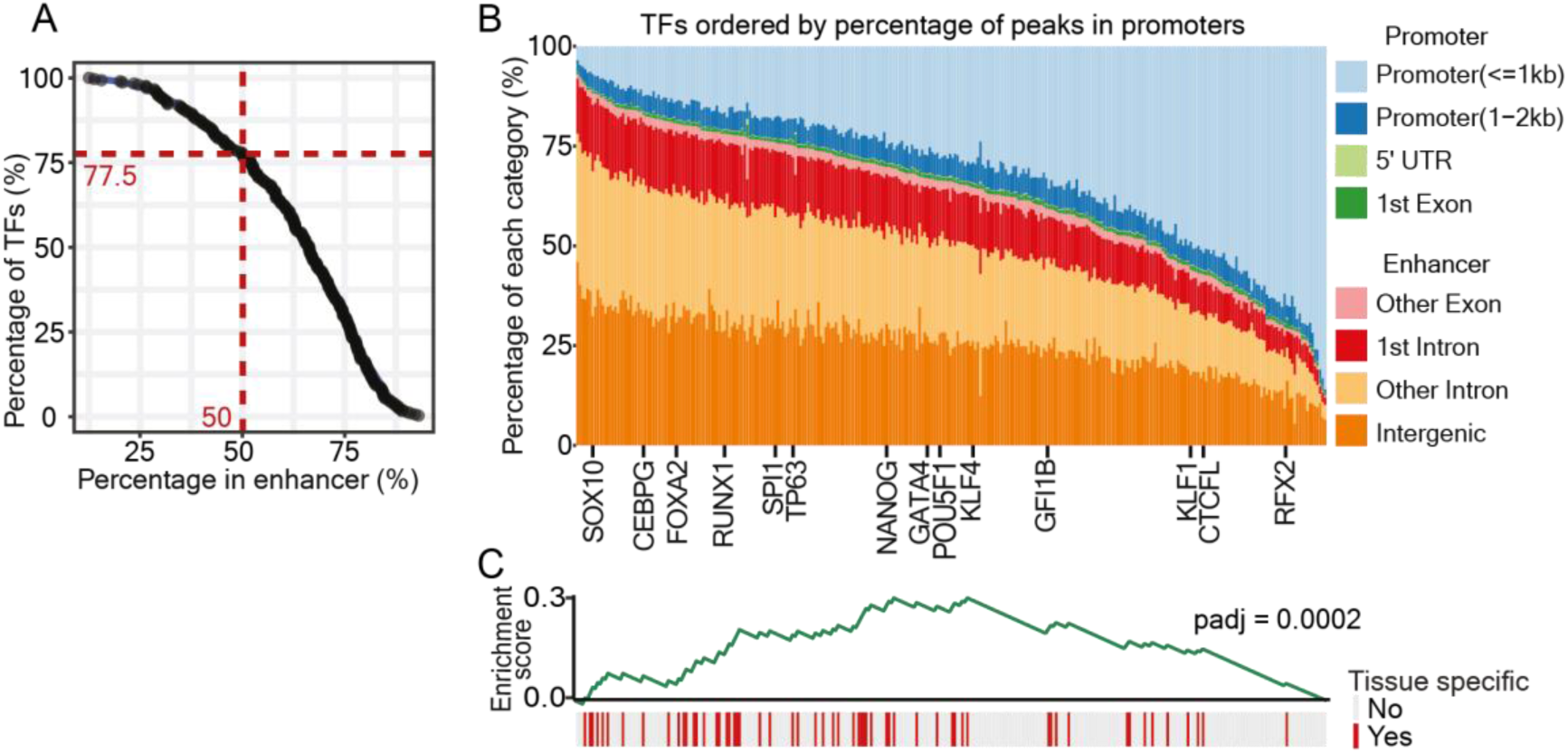
Tissue-specific TFs predominantly bind to enhancers. (A) The percentage of TF binding sites in putative enhancers. The human genome was split into several categories: Promoter (<=1kb), Promoter (1-2kb), 5’ UTR, and 1st Exon, Other Exons, 1st Intron, Other Introns, and Intergenic; these categories were further grouped into a promoter-proximal class (Promoter (<=1kb), Promoter (1-2kb), 5’ UTR, and 1st Exon) and an enhancer class (Other Exons, 1st Intron, Other Introns, and Intergenic). Out of 296 human TFs, 77.5% have at least 50% of their binding sites in the enhancer class of the genome. (B) Genomic location analysis of binding sites of 296 human TFs. The percentage of binding sites of each TF in different categories (as described in A) was calculated, and indicated with different colors. TFs were ordered by the percentage of binding sites within the promoter-proximal class. Several example TFs are marked at the bottom of the figure. (C) Gene Set Enrichment Analysis (GSEA) on tissue-specific TFs and their enhancer binding. The red bars mark the tissue-specific TFs. The order of TFs is consistent with (B). Grey bars represent TFs that do not show tissue-specific gene expression. The GSEA enrichment score is represented by the green line (padj: 2.0e-4).

As expected, we found that the majority of TFs (77.5%) mainly bind in cis-regulatory regions that are distal from the promoter (Figure 1A). These binding sites will not necessarily all be functional, however, they are not close to gene promoters and contain the majority of the enhancers. For the purpose of this study we will refer to them as enhancers. However, different TFs show different binding distributions, with a preference in either the promoter range or in the enhancer range (Figure 1B). Given the relevance of enhancers in cell type-specific gene regulation, we reasoned that cell type-specific TFs would have a larger fraction of peaks in enhancers than constitutively expressed TFs and performed Gene Set Enrichment Analysis (GSEA) (Sergushichev, 2016) on TF expression in different tissues. We defined tissue-specific TFs using previously established categories based on gene expression patterns using RNA-seq from human tissues, including tissue-enriched genes, group-enriched genes, and tissue-enhanced genes (Human Protein Atlas; see methods for details) (Uhlén et al., 2015) (Figure 1C). GSEA showed that TFs mostly binding to enhancers are enriched for tissue-specific expression (adjusted p value = 2.0e-4) (Figure 1C) (Supplementary Table S2). For example, SOX10 is a critical TF during neural crest and peripheral nervous system development (Kim et al., 2014), while TP63 is a master regulator in epithelial development (Soares et al., 2019). Both of these tissue-specific TFs showed a very high percentage of enhancer-binding, 93% for SOX10 and 82% for TP63 (Figure 1A).

Taken together, our analysis of transcription binding sites confirmed that distal cis-regulatory elements are especially relevant for tissue-specific TFs. This emphasizes that including enhancer information in computational methods for predicting key TFs in cell fate determination could be highly beneficial.

### ANANSE: an enhancer network-based method to identify transcription factors in cell fate changes

Starting from the premise that the majority of TFs predominantly bind to enhancer regions, we developed ANANSE, a network-based method that uses properties of enhancers and their GRNs to predict key TFs in cell fate determination (Figure 2). As trans-differentiation is an ideal model for studying cell fate conversions controlled by key TFs, we set out to use this model to validate our computational approach. In the following paragraphs a conceptual overview of ANANSE is provided. Subsequently we will validate each of the steps involved.

**Figure 2.**
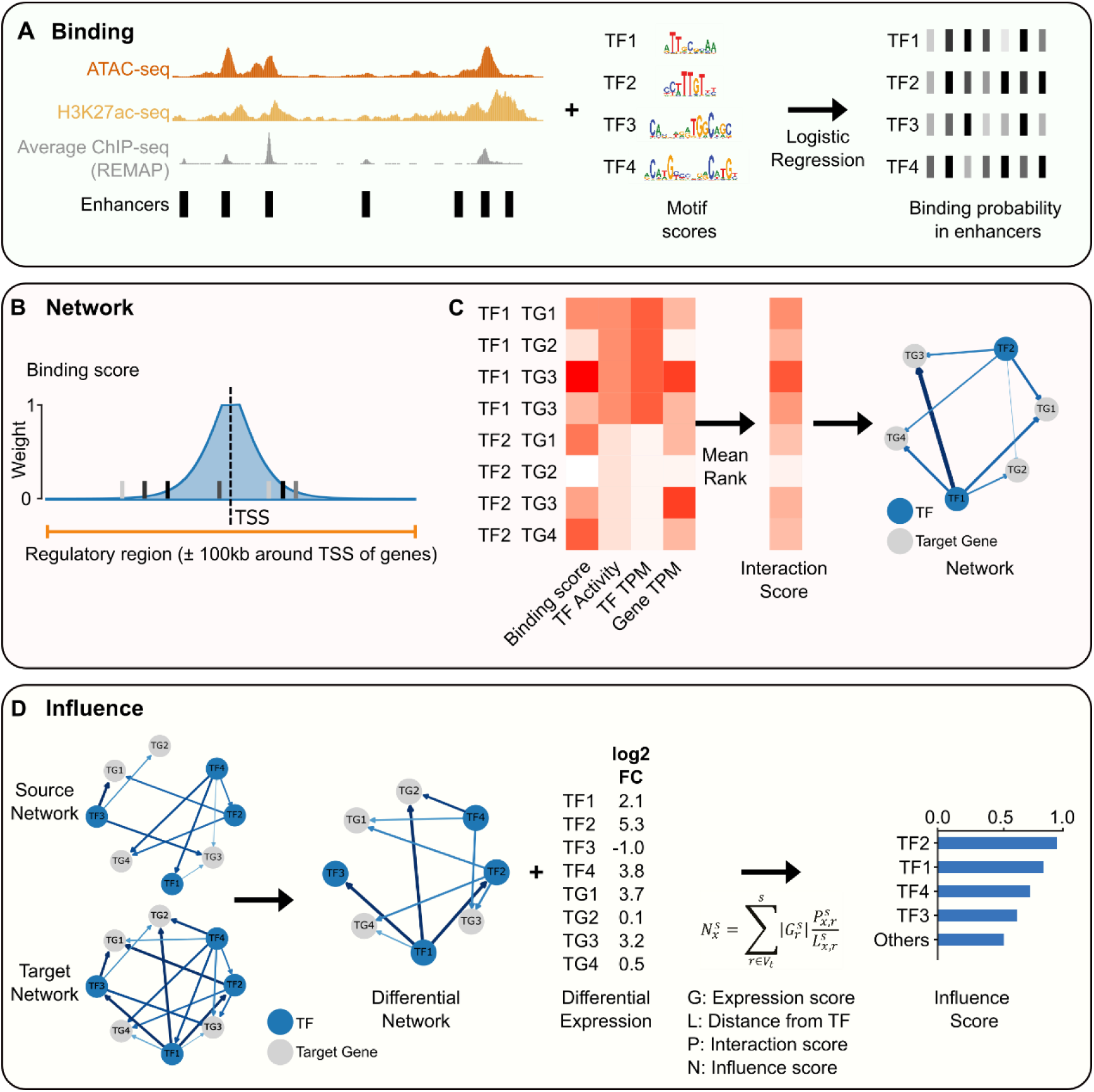
An overview of the ANANSE method. ANANSE consists of three different modules: binding prediction, network inference and influence score calculation. **(A)** TF binding prediction using a supervised model. The binding probability of all TFs with an associated motif is calculated using on the basis of four input data types: 1) a set of reference cis-regulatory regions (here based on REMAP (Chèneby et al., 2017) ChIP-seq integration), 2) genome-wide enhancer activity measurements (ATAC-seq and/or H3K27ac ChIP-seq), 3) the REMAP average ChIP-seq intensity and 4) TF motif scores. The bars in the right panel represent the predicted binding probability of four TFs in the enhancers shown in the left panel. **(B)** A schematic overview of the first step in gene regulatory network inference, calculation of the TF-Gene binding score. A binding score of each TF and gene combination is calculated by aggregation of all enhancers near a gene, weighted by a distance function. The orange line shows 100kb up-and downstream of the TSS of the corresponding target gene, the range that is used to include enhancers for calculation. The bars represent the predicted TF binding probabilities within the 100kb range around the gene. The height of the shaded light blue area represents the weight calculated based on the linear genomic distance from TSS of the target gene to the enhancers (Wang et al., 2016). For example, the distance weight for the distance of 1kb from the TSS is 1, and for the distance of 100kb from the TSS is 0. **(C)** A schematic overview of the gene regulatory network inference using rank aggregation. The heatmap on the left represents the input for each TF and target gene (TG) combination: the binding score (according to **B**), the genome-wide TF activity, the TF expression level (transcripts-per-million; TPM) and the target gene expression level (TPM). All four scores are ranked and scaled from 0 to 1, and the mean of the four scores of each TF-Gene pair is defined as the interaction score (right heatmap) of the corresponding TF-Gene pair. **(D)** Overview of the influence score calculation. The influence score represents how well the expression differences between two cell types can be explained by a TF. First, a differential GRN is calculated between source and target cell type (left). Then, the influence score is calculated based on the gene expression log2 fold change, the distance from the TF to the gene in the predicted network, and the interaction score in the differential network between TF and gene (middle). The barplot (right) shows the ranked influence score of all TFs calculated from the differential GRN.

First, we inferred cell type-specific TF binding profiles for each cell type (Figure 2A). The input data of ANANSE consists of genome-wide measurements of enhancer activity (defined below) and transcription factor motifs. We inferred the TF binding probability based on a supervised model that integrates the enhancer activity combined with TF motif scores.

Second, we constructed cell type-specific GRNs based on the inferred TF binding probability, the transcription factor activity, and the expression levels of the TF and predicted target genes (Figure 2B, 2C). The nodes in the network represent the TFs and their target genes. In this network, a TF node can also be a target gene of another TF. The TF-gene interaction scores, represented by edges of the network, are calculated based on the predicted TF binding probability, the distance between the enhancer and the target gene, the genome-wide TF activity, and the expression of both the TF and the target gene. By integrating these data, ANANSE determines the interaction score of each TF-gene pair.

Third, we calculated the “influence score” (Cahan et al., 2014; Rackham et al., 2016), a measure of importance of a TF in explaining transcriptional differences between two cell types (Figure 2D). In this step, the difference in TF-gene interaction scores (the inferred networks, 2C) between the source and the target cell types is calculated. This differential network is combined with the expression differences between the cell types to determine the influence score.

The details of the algorithms are described in the following sections.

### Transcription factor binding can be predicted by the motif score in combination with the enhancer activity

Sequence-specific TFs bind to their cognate DNA motifs in the genome and activate or repress their target genes. To infer the target genes of a TF, the genomic binding sites of this TF are informative. ChIP-seq has been broadly used to identify TF binding sites at a genome-wide scale. However, it is unfeasible to perform ChIP-seq for every TF in all cell types, e.g. due to the availability and quality of the TF antibodies. Therefore, it would be highly beneficial to be able to predict binding sites of individual TFs in a given cell type.

Here, we used a conceptually simple logistic regression classifier to predict the TF binding probability in putative enhancers based on the TF motif z-score, the enhancer activity and (optionally) the average TF binding signal in these regions (see Methods for details). Our model uses a predefined set of putative enhancers as input. In this work, we used a set of 1.3 million putative enhancer regions based on an integration of all TF ChIP-seq peaks from ReMap 2018 (Chèneby et al., 2017). Alternatively, putative enhancers can be based on genome-wide measurements that provide relatively accurate estimates of enhancer locations, such as ATAC-seq, DNaseI-seq or EP300 ChIP-seq. The enhancer activity is based on two genome-wide assays: chromatin accessibility as measured by Assay for Transposase Accessible Chromatin with high-throughput sequencing (ATAC-seq) (Buenrostro et al., 2013) and the presence of the post-translational histone modification H3K27 acetylation as measured by ChIP-seq.

To train and evaluate our model, we used ChIP-seq peaks of 237 TFs in 6 cell lines (hESC, Hep-G2, HeLa-S3, K562, MCF-7 and GM12878) from REMAP (Chèneby et al., 2017). We examined the locations of the TF peaks by overlapping with our enhancer reference, and only the subset of peaks that overlapped with these enhancer regions was kept. We downloaded and mapped public ATAC-seq and H3K27ac ChIP-seq data for these cell lines (see Supplementary Table S1). For both assays, we determined the number of reads in regions of 200 bp (ATAC-seq) or 2kb (H3K27ac) centered at the enhancer summit. Read counts were log-transformed and quantile normalized. For each enhancer, we scanned for motifs in a 200bp region centered at the peak summit using GimmeMotifs (Bruse and Heeringen, 2018; van Heeringen and Veenstra, 2010). The motif z-score was calculated by GimmeMotifs with the GC%-normalization option. The log-odds score based on the positional frequency matrix is normalized by using the mean and standard deviation of scores of random genomic regions. These random regions are selected to have a similar GC% as the input sequence.

The goal of the binding model in ANANSE is to predict binding for all TFs, based on a supervised model. However, not all TFs have ChIP-seq available for training. Therefore, we implemented a two-pronged approach. We trained a TF-specific supervised model for each TF for which we had training data, but we also trained a general classifier based on all TFs. In this manner, we can use a more performant TF-specific model for TFs that have ChIP-seq training data, but can still predict binding for all other TFs, as long as they have an associated motif. In both cases, the input data for the final trained model is identical, however, the TF-specific models will be better tuned to the binding patterns of their associated TF. To test the prediction performance of our model, we established a stringent cross-validation procedure (Schreiber et al., 2020). For each TF we trained on binding in all enhancers, except those on chromosomes chr1, chr8 and chr23 (held-out chromosomes). In addition, we only included the TFs in the evaluation for which we had more than one cell type available. In turn, each cell type was left out (held-out cell types) and the classifier was trained on data of the other cell type(s). In this manner, performance metrics are calculated based on enhancers located on held-out chromosomes in held-out cell types.

We evaluated the performance of the ANANSE binding model using the AUC (Area Under Curve) of the Precision Recall curve (PR) as well as the (Receiver Operating Characteristic) ROC curve (Figure 3A). For comparison, we included two baselines. The “random” baseline represents the performance that would be observed by “random guessing” (0.5 for the ROC AUC; the proportion of positives in the evaluation set for the PR AUC). The more stringent “Average ChIP-seq” baseline represents the performance that would be observed if the binding is predicted only by the number of different TFs in REMAP that bind to an enhancer (i.e. the predicted binding probability is directly proportional to the number of REMAP peaks overlapping an enhancer). We compared two versions of the ANANSE model, one with the average ChIP-seq peak signal of REMAP included, and one where this is not included (Figure 3A; “With average” and “Without average”, respectively). The model where the average signal is not included is more representative of the performance on other reference enhancer sets, or in other species. The median PR AUC of 0.28 is significantly higher than that of the random baseline (median PR AUC 0.02, p Wilcoxon < 1e-58) as well as the average baseline (median PR AUC 0.18, p Wilcoxon < 1e-38). When we include the average signal, the performance is significantly improved (median PR AUC 0.38, p Wilcoxon < 1e-39). The full ANANSE model also improves on the individual components, as models based on motif scores (median PR AUC 0.06), ATAC-seq (median PR AUC 0.19) or H3K27ac alone (median PR AUC 0.13), while higher than the random baseline, do not significantly outperform the average ChIP-seq baseline. Finally, the model performs well, even when only one of the assays is used (median PR AUC of 0.28 and 0.31 for ATAC-seq and H3K27ac respectively).

**Figure 3.**
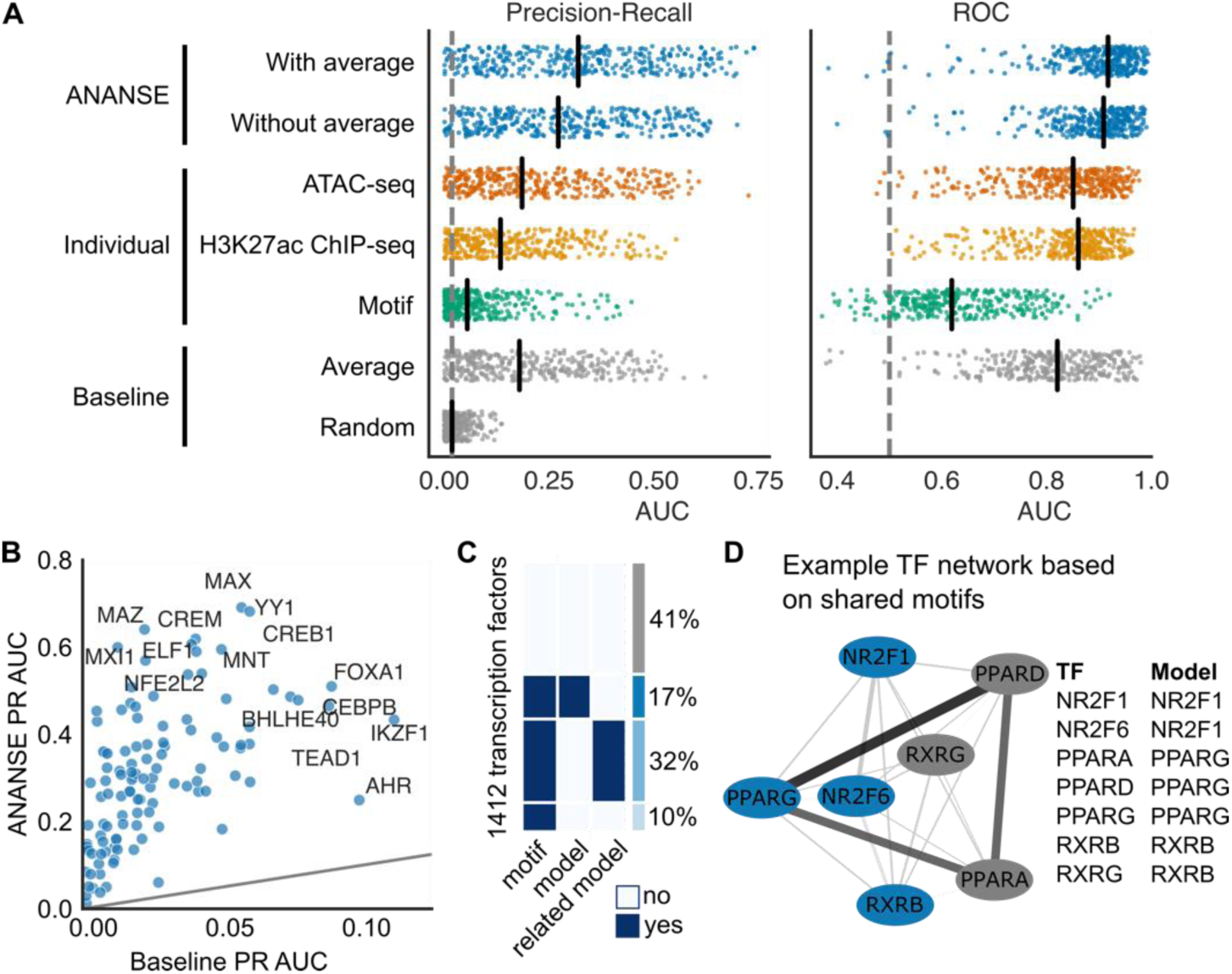
The performance of predicting TF binding sites using TF motif scores and enhancer activities. (A) Evaluation of the TF binding prediction model in ANANSE. Shown is the area under the curve (AUC) of the Precision-Recall (PR; left) and Receiver-operator characteristic (ROC; right) of the prediction performance using REMAP ChIP-seq peaks as a reference. Plotted is the performance of 237 TFs in 6 cell lines. PR AUC and ROC AUC metrics were calculated using cross-validation. The performance is compared to two baselines (grey): random (proportion of positives for PR, 0.5 for ROC) and the average number of REMAP TF ChIP-seq peaks per enhancer. Performance on individual input data types is shown as reference: ATAC-seq (orange), H3K27ac ChIP-seq (yellow) and motif scores (green). Two ANANSE predictions (integration of ATAC-seq, H3K27ac ChIP-seq and motif scores; blue) are compared, based on the inclusion of the average REMAP ChIP-seq signal. Both models perform better than the baselines. (B) The scatterplot shows the improvement of the full ANANSE model compared to the random baseline, with some example factors highlighted. (C) An overview of how many human TFs have an associated model in ANANSE. Out of 1412 TFs, 17% have a trained model available, 32% of TFs have a model trained on a related TF (based on motif similarity, see D) and 10% have a motif and can use the general model. The remaining 41% of TFs do not have an associated motif. (D) An example of determining related TFs for model sharing. The network illustrates the similarity between a selection of nuclear receptors, as determined by the Jaccard index of their associated motifs (edge color and size). There is no trained model for PPARD, but it shares many motifs with PPARG, so it uses the PPARG model weights with PPARD motif scores.

For comparison to methods that train more complex supervised transcription factor-specific models, we also validated our model on the validation chromosomes (chr1, chr8, and chr21) in the validation cell types of the ENCODE-DREAM transcription factor binding challenge (ENCODE-DREAM, 2017). As a comparison, we used the Virtual ChIP-seq predictions (Karimzadeh and Hoffman, 2018). This is a newly developed supervised artificial neural network method to predict individual TF binding, which shows comparable results compared to the top ENCODE-DREAM entries. In this evaluation (Supplementary Figure S1A and B), our model scores better for some factors, such as CEBPA in liver and MAX in liver and K562 cells, while Virtual ChIP-seq performs better for other factors, most notably CTCF. This illustrates that the binding prediction of ANANSE is comparable to state-of-the-art approaches. A caveat here is that other methods, such as Virtual ChIP-seq, predict binding genome-wide, while ANANSE needs a set of putative enhancers as input. An advantage of the relatively simple model of ANANSE is that it generalizes to other TFs and other species, which will not have the abundance of training data provided by ENCODE for mouse and human. As another evaluation, we compared the ANANSE binding predictions to predictions based on DNase I footprinting (Supplementary Figure S1C and D) (Vierstra et al., 2020). Compared to DNase I footprints, ANANSE predictions have higher recall at the same precision (median 0.78 vs 0.01).

In total, these analyses illustrated that we established a TF binding site prediction method, which can be applied on the basis of one or two experimental measurements (ATAC-seq and/or H3K27ac ChIP-seq) and which shows state-of-the-art performance in prediction of TF binding.

### ANANSE predicts cell type-specific gene regulatory networks

Using the inferred cell type-specific binding profiles, we sought to determine the interactions of TFs and their target genes (TF-gene) to establish cell type-specific GRNs, represented by the TF-gene interaction score. To calculate these scores, we first identified all enhancers for each target gene and their associated binding scores. In our TF binding prediction model, we used H3K27ac ChIP-seq and ATAC-seq as training data. For each gene, we took all enhancers that are located within a maximum distance of 100 kb of the transcription start site (TSS). Subsequently, the strength of a TF-gene interaction in the network was defined by the sum of the predicted TF binding strength in all identified enhancers of the target gene weighted by the distance (Figure 2B), similar to the regulatory potential (Wang et al., 2016). The distance weight was calculated from the linear genomic distance between the enhancer and the TSS of a gene, such that distal enhancers receive a low weight and nearby enhancers have a high weight (Figure 2B). This model resulted in a TF-gene binding score, indicating the TF-target gene binding intensity for all combinations of TF and target gene pairs.

Next to the incorporation of the genome-wide measurements of enhancer activity via the TF-Gene binding score, we also calculated a measure of TF activity directly from the genome-wide enhancer activity. Here, we used a method similar to the motif activity response analysis (Balwierz et al., 2014; Suzuki et al., 2009) as implemented in GimmeMotifs (Bruse and Heeringen, 2018). In this approach, the enhancer activity (ATAC-seq and H3K27ac signal) is modeled as a linear function of motif scores using penalized regression. The coefficients of the motif scores can be interpreted as an estimate of motif activity. For all TFs we used the maximum activity of all associated motifs as TF activity.

Finally, based on the assumption that the interaction of every TF-gene pair in a specific cell type is proportional to their relative expression, we included the expression level of the TF and the target gene, the TF and target expression scores. We ranked the expression level of the TF and the target gene, initially expressed as transcripts per million (TPM) within each cell type to a normalized expression between 0 and 1, with the lowest expression as 0 and highest as 1.

To calculate the TF-gene interaction score, we combined the TF-gene binding score, the TF activity, and the TF and target expression scores using mean rank aggregation (Figure 4C). This score represents the strength of the regulatory interaction between a TF and a target gene. In this approach, a “target gene” can also be a TF gene; a TF-gene interaction can represent a TF regulating the expression of a TF gene. Together, all TF-gene interaction scores represent a cell type-specific GRN.

**Figure 4.**
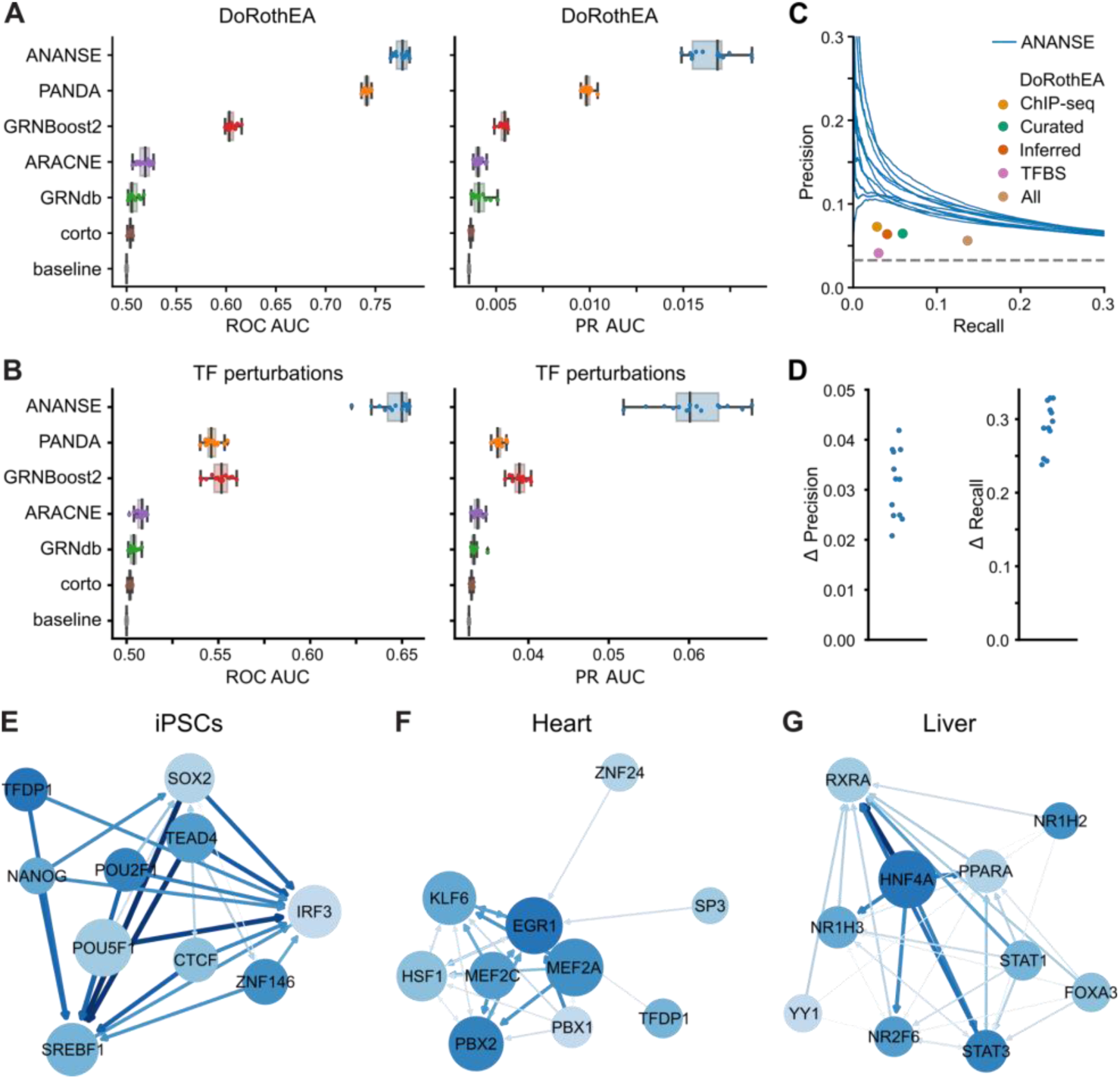
Prediction of tissue-specific enhancer gene regulatory networks. **(A)** Evaluation of the predicted networks using the curated interactions of the DoRothEA database. The left panel shows the AUC of ROC for 15 different tissues in a boxplot, with individual tissues marked as dots. The right panel shows the PR AUC. The ANANSE predicted networks (blue) are compared to other GRN inference approaches trained on GTEx expression data for the same tissues (PANDA in orange, GRNBoost2 in red; ARACNE in purple and corto in brown) and on the GRNdb networks inferred using SCENIC on single cell RNA-seq data from the same tissues (green). The random baseline is shown in gray. **(B)** The same evaluation as in A) using a reference of differentially expressed genes after TF perturbation. **(C)** Comparison of the tissue-specific GRNs inferred by ANANSE to the different types of interactions in DoRothEA, using the TF perturbations as a reference. The PR curves of the inferred networks for the different tissues are plotted (ANANSE; blue), and the different interactions present in DoRothEA are represented by dots: interactions predicted by TF ChIP-seq binding near genes (orange), curated interactions from the literature (green), interactions inferred using ARACNe-VIPER (red) and interactions predicted by TF motif scores in the promoter (purple). The union of all DoRothEA interactions is shown in brown. **(D)** The difference of the ANANSE GRNs with the union of DoRothEA interactions in C) expressed as the difference in precision at the same recall (left panel) and the difference in recall at the same precision (right panel). **(E)** Example network predicted for iPSCs. The blue circles show the top 10 TFs in this cell type, ranked by the outdegree in the top 100,000 edges. The size of the circle indicates the target gene number of the corresponding TF. The size and color of the blue arrows are relative to the interaction score between the two TFs. The color of the circle indicates the expression level of the corresponding TF. **(F)** Example network predicted for heart, visualized as in E). **(G)** Example network predicted for liver, visualized as in E).

To evaluate the quality of the GRNs inferred by ANANSE, we created GRNs for 15 tissues: adrenal gland, brain, colon, esophagus, heart, liver, lung, ovary, pancreas, prostate, skeletal muscle, skin, small intestine, spleen, and stomach. We collected ATAC-seq, H3K27ac and RNA-seq from public repositories (see Supplementary Table S1) and predicted binding profiles and GRNs using ANANSE. As comparison, we included five other GRN inference methods. We downloaded ARACNE, corto and PANDA networks created using GTEx expression data (Glass et al., 2013; Margolin et al., 2006; Mercatelli et al., 2020). We downloaded the GTEx expression data and created GRNs using GRNBoost2 (Moerman et al., 2019). We downloaded GRNs created by SCENIC (Aibar et al., 2017) with tissue single cell data as input from GRNdb (Fang et al., 2021).

To provide a comprehensive overview of GRN quality, we used four different types of reference datasets to calculate performance metrics: 1) regulatory interaction databases containing known TF-target gene interactions, 2) differential expression measurements after TF perturbation, 3) gene co-expression data and 4) Gene Ontology (GO) annotation (The Gene Ontology, 2019). We obtained the TF-gene interactions from four databases of regulatory interactions, DoRothEA (Holland et al., 2020), RegNetwork (Liu et al., 2015), TRRUST (Han et al., 2017) and MSigDB C3 (Liberzon et al., 2011). DoRothEA is a gene set resource network containing different types of TF and target interactions. For this comparison, we only used the literature-curated interactions. RegNetwork is an integrated database of transcriptional and post-transcriptional regulatory networks in human and mouse, TRRUST is an expanded reference database of human and mouse transcriptional regulatory interactions and MSigDB C3 is a collection of gene sets that represent potential targets of regulation by TFs. While the TF-target gene databases are usually curated, they contain relatively few interactions. As a source of a more genome-wide validation we downloaded the “TF_Perturbations_Followed_by_Expression” data set from Enrichr (Chen et al., 2013; Kuleshov et al., 2016). This is a curated collection of genes that significantly change expression after TF perturbation. We downloaded co-expression data for human genes from the COXPRESdb database (Obayashi et al., 2019). All TF-gene pairs with either a correlation >= 0.6 or >= 0.8 were used as true positives. Finally, we used TF-gene pairs that were annotated with at least one common GO term as true positives.

To compare with the tissue-specific GRNs inferred by other methods, we selected per reference the Cartesian product of TFs and target genes of the sets of TFs and target genes in the reference. If a specific interaction was not present in the inferred GRN, we used the minimum interaction score. Supplementary Note S1 contains more details on the benchmark procedure. We evaluated the GRNs by calculating the PR AUC and ROC AUC, as compared to the reference interactions (Figure 4A and B; Supplementary Figure S2). For the ANANSE networks, the median AUC ranges from 0.61 using the RegNetwork reference (Supplementary Figure S2A) to 0.77 using DoRothEA (Figure 4A), while the median AUC of randomized networks is close to 0.5. When we compared ANANSE with other published GRN inference methods, all five methods show a significantly lower AUC using DoRothEA (Figure 4A) or TF perturbation references (Figure 4B). This holds true for all other references, except for the correlation reference, where PANDA scores higher (Supplementary Figure S2A). Some of the reference databases contain very few interactions (the positives in this evaluation) as compared to all possible interactions (which determine the negatives). For instance, the fraction of positive interactions is 0.09% in TRRUST, 0.07% in RegNetwork, and 0.33% in MSigDB C3. Therefore, we also evaluated the predicted networks using the PR AUC (Figure 4D and Supplementary Figure S2B). In absolute terms, the PR AUC is considerably lower than the ROC AUC, especially for the DoRothEA, TF perturbation, MSigDB C3, TRRUST, and RegNetwork reference sets (median PR AUC of 0.0168, 0.0601, 0.0297, 0.0123, and 0.0187, respectively), but for all tissues there is a relatively large and statistically significant difference between the predicted GRN and the randomized network (p Wilcoxon = 9.86e-22) (Figure 4A and B, Supplementary Figure S2B). The expression-based networks of the other five published methods show significant lower mean PR AUC when compared with ANANSE for all seven different types of reference datasets (Figure 4A and B and Supplementary Figure S2B).

To further evaluate the ANANSE networks, we compared them to all different types of interactions present in DoRothEA, using the TF perturbations as a reference. In Figure 4C the PR curves of the inferred networks for the different tissues are plotted, and the different interactions present in DoRothEA are represented by dots. These include interactions predicted by TF ChIP-seq binding near genes (ChIP-seq), curated interactions from the literature (Curated), interactions inferred using ARACNe-VIPER (Inferred) and interactions predicted by TF motif scores in the promoter (TFBS). In addition the union of all sets is shown (All). The difference in recall at the same precision, and precision at the same recall between the ANANSE networks and the union set (All) of DoRothEA is shown in Figure 4D. These results illustrate that, using this benchmark, ANANSE-inferred networks better predict genes deregulated by TF perturbation.

The inclusion of enhancer information in the GRN inference is one of the unique features of ANANSE. To test if incorporating enhancer activity indeed leads to an improved performance, we compared the enhancer-based approach of GRN with an expression-based model and two promoter-based models (Supplementary Figure S3). Overall we found that the incorporation of enhancer information leads to better network inference, as tested by these benchmarks. The choice of distance and the method of combining enhancers has a smaller effect, with a 100kb distance performing marginally better than 250kb and the distance-weighted sum performing better than the mean or sum with a uniform weight (Supplementary Figure S4).

To qualitatively assess the cell type-specific GRNs predicted by ANANSE, we chose one well-studied cell type (iPSC) and two tissues (heart and liver) and constructed their GRNs using the top ten predicted TFs of each cell type, as ranked by outdegree. The GRN of iPSCs includes well-known pluripotency factors such as POU5F1, NANOG and SOX2 (Figure 4E). The GRNs of heart and liver tissues contain marker genes, such as the myocyte factors MEF2A and MEF2C in heart (Figure 4F), and HNF4A and FOXA3 in liver (Figure 4G).

Taken together, our benchmarks and examples demonstrate that GRNs generated by ANANSE allow for meaningful cell type-specific prioritization of TFs.

### ANANSE accurately predicts key transcription factors for trans-differentiation

Having established that ANANSE-inferred GRNs can enrich for biologically relevant regulatory interactions, we aimed to use these GRNs to identify key TFs that regulate cell fate determination. To this end, trans-differentiation is a good model, as experimentally validated TFs have been determined for various trans-differentiation strategies. Here, we first inferred the GRNs for all cell types using our ANANSE approach. The ANANSE-inferred GRN difference between a *source* and a *target* cell type, was calculated to represent the differential GRN between two cell types, which contains the GRN interactions that are specific for or higher in the target cell type. Subsequently, using an approach inspired by Mogrify (Rackham et al., 2016), we calculated the influence score of TFs for these trans-differentiations by determining the differential expression score of its targets weighted by the regulatory distance (see Methods for details).

To evaluate the prediction by ANANSE, we used experimentally validated TFs for several trans-differentiation strategies. For this, we collected TFs for seven trans-differentiation strategies with fibroblasts as the source cell type. The target cell types include astrocytes (Caiazzo et al., 2015), cardiomyocytes (Fu et al., 2013; Ifkovits et al., 2014), hepatocytes (Nakamori et al., 2017; Simeonov and Uppal, 2014), iPSCs (Huangfu et al., 2008; Takahashi et al., 2007; Yu et al., 2007), keratinocytes (Kurita et al., 2018), macrophages (Feng et al., 2008; Xie et al., 2004), and osteocytes (Li et al., 2017; Yamamoto et al., 2015) (Table 1). We used ATAC-seq and H3K27ac ChIP-seq data of these cell types to create cell type-specific GRNs, and then calculated TF influence scores and ranked the TFs in each cell type.

**Table 1.**
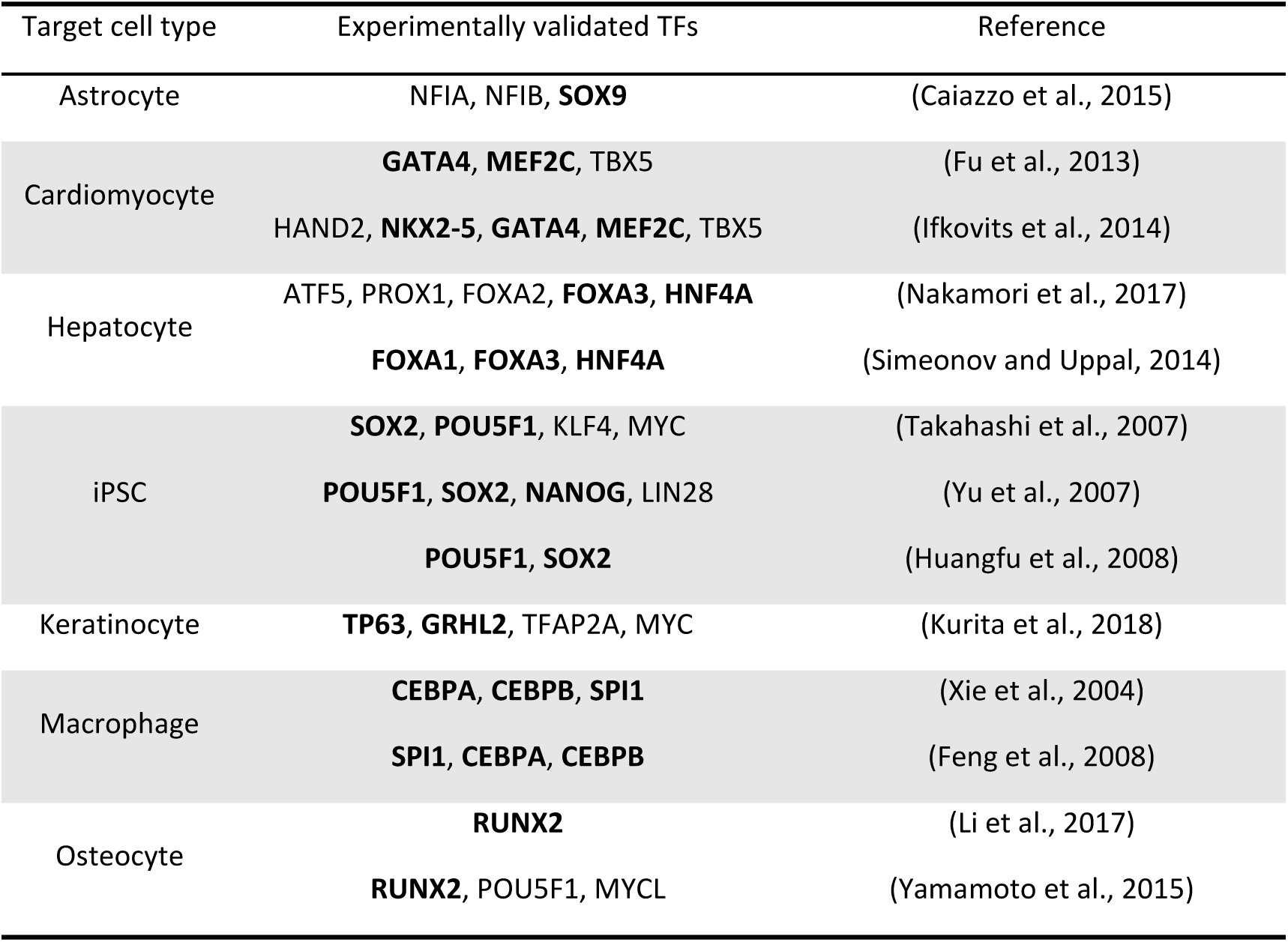
The summary of seven experimentally validated trans-differentiations from fibroblast to target cell types. Experimentally validated TFs that were identified by ANANSE are highlighted in bold.

When we calculate TF influence scores from cell type-specific GRNs, it is important to decide what size of GRN should be chosen in terms of the top number of edges. We inferred the key TFs for the seven trans-differentiations using six different sizes of GRNs (10K, 50K, 100K, 200K, 500K, and 1M edges; Supplementary Figures S4,5). These results show that the approach is relatively invariant to the GRN size, with performance starting to decrease at 1 million edges. Here we chose a GRN size of 100K interactions for all following analyses.

Using GRNs with 100K edges, we predicted the top 10 TFs for all seven trans-differentiations. In all cases, many of the experimentally defined TFs are included in the top 10 factors predicted by ANANSE (Table 1). For example, ANANSE predicts CEBPA, CEBPB and SPI1 for reprogramming fibroblasts to macrophages (Feng et al., 2008; Xie et al., 2004) and HNF4A, FOXA1 and FOXA3 for reprogramming to hepatocytes, which are consistent with the experimental trans-differentiation strategies (Nakamori et al., 2017; Simeonov and Uppal, 2014).

To evaluate if the inclusion of enhancer information in the GRN inference in ANANSE resulted in more accurate predictions of TFs for trans-differentiation, we compared the enhancer-based approach of ANANSE with a promoter-based model, which does not take into account promoter-distal regulatory elements, and with a model based on only gene expression data (Figure 5A). We created both expression and promoter based GRNs of the seven source and target cell type combinations. For expression-based GRNs, we used only the mean of the scaled TPM of TFs and genes together as the interaction score of TFs and genes. For the promoter-based GRNs, we selected the highest binding score of TFs within 2kb of the TSS of the corresponding gene as the binding score of the TF-gene pair. Subsequently, the mean of the scaled TPM of the TF and the gene together with the binding score determines the interaction score of the TF and gene (Figure 4B). We then inferred the key TFs for the seven trans-differentiations using ANANSE and these two types of GRNs. The ANANSE influence score based on the enhancer GRNs includes 53% of the known TFs in the top four predictions (Figure 5A and Supplementary Table S6). In contrast, using the influence score based on the promoter GRN or the expression GRN, we could recover only 5% and 14% of the known TFs in the top four predictions (Figure 5A, Supplementary Figures S7 and S8A and Supplementary Table S6). These results demonstrate that using enhancers in the construction of GRNs significantly improves the prediction of relevant TFs in cell fate determination.

**Figure 5.**
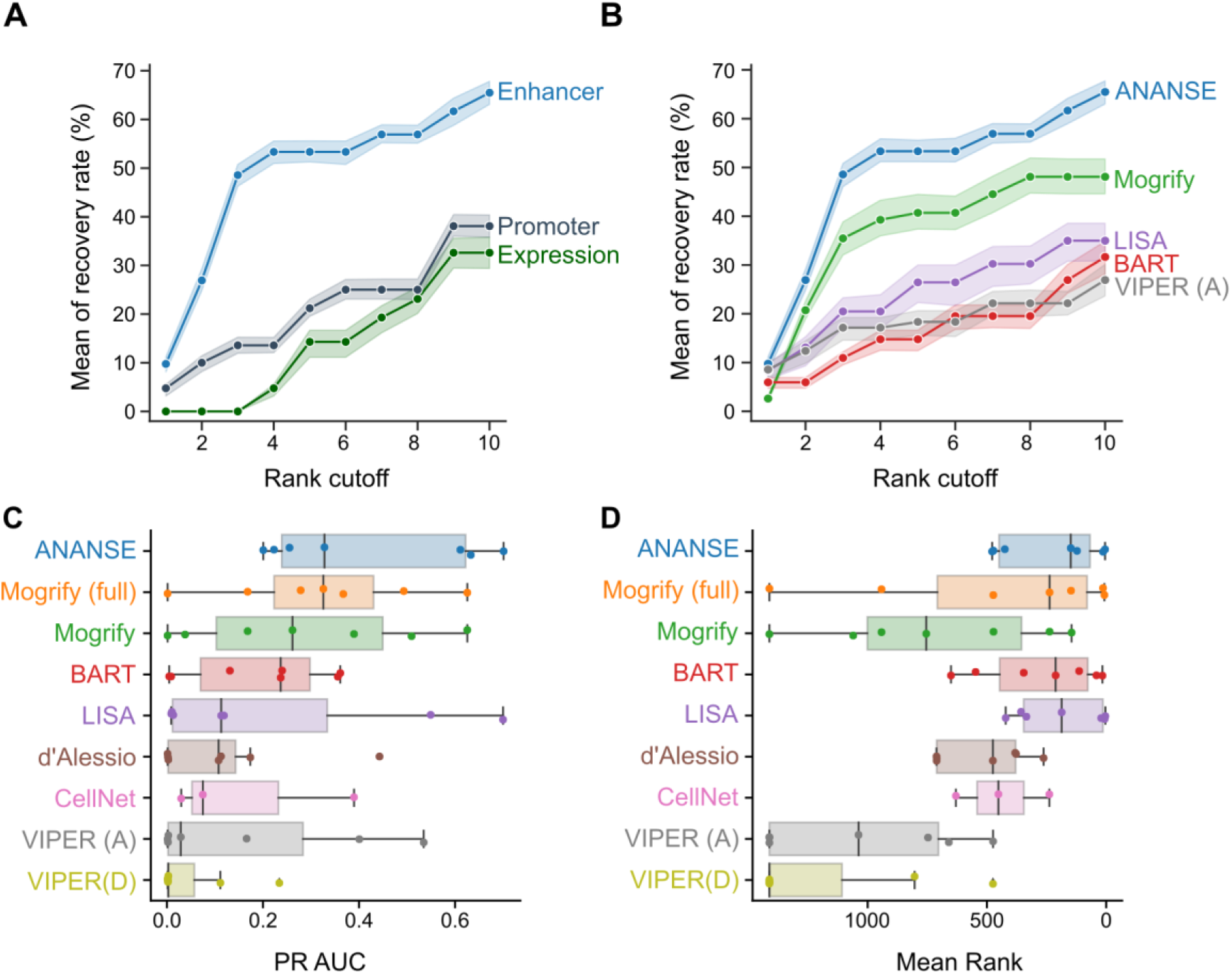
Evaluation of the performance of ANANSE using experimentally validated trans-differentiation strategies. **(A)** The line plots show the comparison of the predicted top TFs for trans-differentiation from cell type-specific networks. Based on the difference between two networks, TFs were prioritized using the influence score calculation implemented in ANANSE. Shown is the fraction of predicted TFs compared to all known TFs based on trans-differentiation protocols described in the literature (y-axis) as a function of the top number of TFs selected (x-axis). The mean of recovery rate is the average of all TF sets when the corresponding trans-differentiation has several different experimental validated TF sets. The shaded area represents the minimum and maximum percentage of corresponding recovered TFs when using six out of seven trans-differentiations. Three different types of networks were used: gene expression (deep green), promoter-based TF binding in combination with expression (deep blue), and enhancer-based TF binding in combination with expression (blue). **(B)** The line plots show the comparison of the predicted top TFs for trans-differentiation based on different computational methods. The y-axis indicates the percentage of experimentally validated cell TFs that are recovered as a function of the number of top predictions, similar as in A). Six different methods are shown: ANANSE (blue), Mogrify (green), LISA (purple), BART (red), and VIPER with ANANSE networks (gray). The shaded area represents the minimum and maximum percentage of corresponding recovered TFs when using six out of seven trans-differentiations. Mogrify and CellNet only contain the top 8 predicted factors. For visualization purposes, only a subset of the evaluated methods is shown. The remaining methods are shown in Supplementary Figure S8B. **(C)** The PR AUC of the same trans-differentiations as in A and B, shown as a boxplot. Individual trans-differentiations are shown as dots**. (D)** The mean rank of the experimentally determined factors of the same trans-differentiations as in A and B, shown as a boxplot. Individual trans-differentiations are shown as dots.

Next, we further quantified the performance difference between ANANSE and previously reported methods, namely Mogrify, LISA, BART, VIPER, CellNet, and the method of D’Alessio et al (Alvarez et al., 2016; Cahan et al., 2014; D’Alessio et al., 2015; Qin et al., 2020; Rackham et al., 2016; Wang et al., 2018) (Figure 5B and Supplementary Figure S8). For Mogrify, we downloaded both the prioritized list of TFs based on TF expression in source cell types and GRN overlap, as well as the full unfiltered list of TFs. For VIPER, we predicted TFs with the DoRothEA network, “VIPER (D)”, and with the GRNs inferred by ANANSE, “VIPER (A)”. For these comparisons, we aimed to include all seven trans-differentiation strategies. In some cases, as data for the exact cell type is unavailable, similar cell or tissue types were used as surrogates. For example, the osteoblast-Sciencell was used to substitute for osteoblast. For CellNet, we used the previously described results of three cell types: hepatocytes, iPSCs and macrophages (Rackham et al., 2016). For LISA, we used all differentially expressed genes from fibroblast to each target cell type as input (Qin et al., 2020). For all methods, the resulting TFs were ranked according to the relevant output score of the method. Using the seven cell type conversions as a reference, ANANSE has the highest recovery at all rank cutoffs up to 10 (Figure 5B and Supplementary Figure S6). ANANSE predicts a mean of 53% TFs using the top four TFs ranked by influence score, while other methods predict a maximum of 39% of TFs with this rank cutoff (Figure 5B and Supplementary Figure S6). When the number of predicted TFs was increased to ten, ANANSE could increase its recovery rate to 65%, while the maximum mean recovery of other methods is 48% (Figure 5B and Supplementary Figure S8). In addition to the mean recovery rate, we also evaluated the PR AUC (Figure 5C) and the mean rank of all known trans-differentiation factors (Figure 5D). In these analyses ANANSE shows the highest median score (PR AUC or mean rank). However, other methods perform nearly as well in these benchmarks, such as Mogrify (both with PR AUC and mean rank) and BART and LISA (mean rank). In summary, these analyses show that including enhancers in the GRN construction significantly improves the prediction of TFs in cell fate conversion and that ANANSE outperforms other established methods, based on experimentally validated trans-differentiation TFs. Our results demonstrate that ANANSE can prioritize biologically relevant TFs in cell fate determination.

### ANANSE identified an atlas of key transcription factors in normal human tissues

The gene expression programs that drive the cellular differentiation programs of different tissues are largely controlled by TFs. To find out which key TFs drive cell fate determination in different tissues, we applied ANANSE to a much wider range of human tissue data. We downloaded H3K27ac ChIP-seq data of 18 human tissues from the ENCODE project (Encode Project Consortium, 2012) and the RNA-seq data of corresponding tissues from the Human Protein Atlas project (Uhlén et al., 2015). Using these enhancer and gene expression data, we constructed tissue-specific GRNs using ANANSE, and then calculated the TF influence scores for each of the tissues when taking the combination of all other tissues as the source tissue (Supplementary table S7). We clustered the 18 tissues based on the correlation between TF influence scores using hierarchical clustering, showing that the influence score captures regulatory similarities and differences between tissues (Figure 6A and Supplementary Figure S). For example, the esophagus and the skin cluster together, as these tissues are composed mostly of stratified squamous epithelial cells, and skeletal muscle and heart tissue are clustered together as both tissues contain striated muscle tissues.

**Figure 6.**
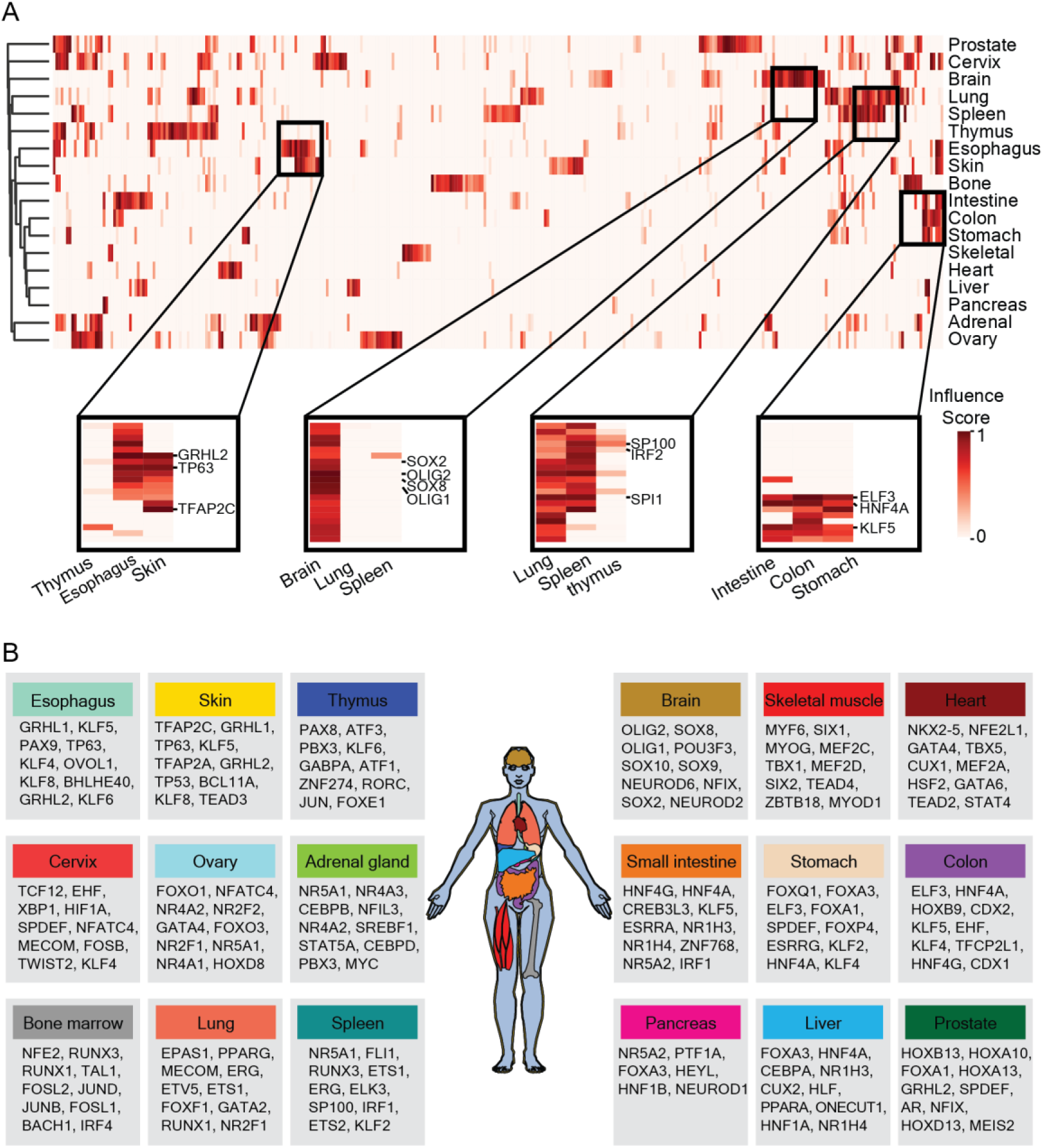
Applying ANANSE to expression data of human tissues to identify key transcription factors. (A) Heatmap of the predicted influence scores of all TFs using ANANASE on data from 18 human tissues. The color in the heatmap indicates the relative influence score, from low to high. The four small heatmaps highlighted below show important TFs in related tissues. (B) The top 10 key TFs of 18 tissues inferred by ANANSE. The color of the tissue is consistent with the tissue name in the box. The order of TF of each tissue is based on the influence score of the TF ranked from high to low.

For all studied tissues, we have provided a rich resource of key TFs of each tissue, with a list of top ten key TFs (Figure 6B). Many TFs in this list are known to play important functions for specific tissues, e.g. ELF3 and KLF5 for stomach, colon, and small intestine (Jedlicka et al., 2008; Katz et al., 2002); TFAP2A, TFAP2C, TP63, and GRHL2 for the skin and esophagus (Dollé, 2009; Qu et al., 2018; Wilanowski et al., 2008); SOX2, SOX8 and OLIG1/2 for brain (Bani-Yaghoub et al., 2006; Meijer et al., 2012; Muto et al., 2009); and SPI1 for lung, spleen and bone marrow (Ohteki et al., 2001)(Figure 6A).

In summary, using ANANSE, we predicted key TFs for 18 human normal tissues. Many of these predicted TFs correlate well with the known literature of these tissues. In addition, the predicted key TFs in each tissue also provide us a rich resource to unveil novel TFs in specific tissues.

## Discussion

Lineage specification and cell fate determination are critical processes during development. They are necessary to form the diversity of cell types that are organized into organs and tissues. TFs form a central component in the regulatory networks that control lineage choice and differentiation. Indeed, cell fate can be switched *in vitro* through manipulation of TF expression (Caiazzo et al., 2015; Feng et al., 2008; Fu et al., 2013; Huangfu et al., 2008; Ifkovits et al., 2014; Kurita et al., 2018; Li et al., 2017; Nakamori et al., 2017; Simeonov and Uppal, 2014; Takahashi et al., 2007; Xie et al., 2004; Yamamoto et al., 2015; Yu et al., 2007). However, the regulatory factors that determine cell identity remain unknown for many cell types. To address this issue, we developed ANANSE, a new computational method to predict the key TFs that regulate cellular fate changes.

We establish TF binding networks for each cell type by leveraging genome-wide, cell type-specific enhancer signals from ATAC-seq, H3K27ac ChIP-seq and TF motif data. ANANSE takes a two-step approach. First, TF binding is imputed for all enhancers using a simple supervised logistic classifier. In contrast to existing methods that aim to predict binding by training TF-specific models (Batsis et al., 2019; Keilwagen et al., 2019; Li et al., 2019a; Quang and Xie, 2019), we also used a more general model that can be applied to all TFs. Logically, our simple model will not be as accurate as complex models, trained for specific TFs. Indeed, the PR AUC and ROC AUC of the ANANSE binding model are lower for a factor such as CTCF than the current state-of-the-art in supervised prediction, as illustrated by the comparison with Virtual ChIP-seq (Supplementary Figure S1; Karimzadeh and Hoffman, 2018). However, the advantage of our model is that it can predict binding for every TF as long as its motif is known. In addition, it can be used for factors for which there is no training data available, and it can also be applied to non-model organisms that lack comprehensive ChIP-seq assays. Our benchmarks show that it performs significantly better than using the motif score alone or enhancer activity alone and that it outperformed a more stringent average ChIP-seq baseline.

Second, we summarized the imputed TF signals per gene, using a distance-weighted decay function (Wang et al., 2016), and combined this measure with TF activity and TF and target gene expression to infer cell type-specific GRNs. In general, there is a lack of gold standards to evaluate cell type-specific GRNs. We used multiple orthogonal types of benchmarks: databases of known, experimentally identified TF-gene interactions, gene expression after TF perturbations, and functional enrichment using Gene Ontology annotation. The databases with known interactions that we used contain only a fraction of true regulatory interactions, and therefore this benchmark is affected by a large fraction of false negatives. All our benchmark evaluations demonstrate that ANANSE significantly enriches for true regulatory interactions. However, it also highlights that GRN inference is far from a solved problem. The PR-AUC values are low, as is generally the case in eukaryotic GRN inference (Chen and Mar, 2018). Our comparison with several established gene expression-based GRN approaches, such as PANDA (Glass et al., 2013), GRNBoost2 (similar to GENIE3) (Huynh-Thu et al., 2010; Moerman et al., 2019) and ARACNE (Margolin et al., 2006) shows that these methods also result in low PR-AUC values. In addition to the improved performance, ANANSE has another clear benefit. While most GRN inference methods need a large collection of samples, ANANSE uses only two or three genome-wide measurements as input: gene expression and enhancer activity (H3K27ac ChIP-seq and/or ATAC-seq).

In contrast to previous approaches, our method takes advantage of TF binding in enhancers, instead of only gene expression differences or TF binding to proximal promoters. This resulted in significantly improved performance, as benchmarked for the GRN inference, as well as on experimentally validated trans-differentiation protocols. It has been previously shown that cell type-specific regulation is much better captured by enhancers as compared to promoter-proximal regulatory elements. For instance, TF binding and chromatin accessibility in distal elements better reflect the cell type identity of hematopoietic lineages than in promoters (Corces et al., 2016; Heinz et al., 2010). Many important transcriptional regulators mainly bind at regulatory regions that are not proximal to the promoter. Indeed, our analysis of the genomic binding distribution of ∼300 human TFs showed that cell type-specific TFs bind in enhancer regions more often than TFs that are more widely expressed (Figure 1C). Therefore, we reasoned that TF binding at enhancers would be essential to model cell fate and lineage decisions. We tested the application of the networks inferred by ANANSE to human *in vitro* trans-differentiation approaches. Earlier work showed that computational algorithms allow characterization of cellular fate transitions and rational prioritization of TF candidates for trans-differentiation (Cahan et al., 2014; Morris et al., 2014; Rackham et al., 2016). We implemented a network-based approach to prioritize TFs that determine cell fate changes. Using a collection of known, experimentally validated trans-differentiation protocols, we demonstrated that ANANSE consistently outperforms other published approaches. This means that cellular trajectories can be characterized using ANANSE to identify the TFs that are involved in cell fate changes. In comparison with a promoter-based approach, we show that using enhancer-based regulatory information contributes significantly to this increased performance (Figure 5A). One noticeable example is the trans-differentiation from fibroblasts or mesenchymal cells to keratinocytes. In current experimentally validated trans-differentiation methods, the epithelial master regulator TP63 is essential for establishing the keratinocyte cell fate (Chen et al., 2014; Kurita et al., 2018). However, TP63 was not predicted in most of the previously published computational methods (Cahan et al., 2014; Morris et al., 2014; Rackham et al., 2016). One plausible explanation is that TP63 is a TF for specific epithelial cells and tissues and it binds predominantly (87%) to enhancers (Andersson et al., 2014; Bulger and Groudine, 2011; Qu et al., 2018; Spitz and Furlong, 2012), whereas previous computational tools do not take enhancer properties into consideration.

We used ANANSE to identify tissue-specific TFs for different human tissues. We predicted the top 10 key TFs for all studied tissues. Many TFs in this list are known for important functions in these specific tissues. For example, some NK homeodomain, GATA, and T-box TFs are found in normal cardiac development, which have important functions during heart specification, patterning, and differentiation (Bruneau, 2013; Kathiriya et al., 2015; Stefanovic and Christoffels, 2015). Many TFS of the SOX family are known to be critical for neural system development in brain tissue (Bani-Yaghoub et al., 2006; Muto et al., 2009). The gastrointestinal tract tissues share a number of high influence score TFs such as ELF3, KLF5, and HNF4A, which play roles in stomach, colon, and small intestine development, and are consistent with the current research on gastrointestinal tract tissues (Figure 6A) (Johnston et al., 2019; Nandan et al., 2014; Thompson et al., 2018). ELF3 is important in intestinal morphogenesis, homeostasis, and disease (Johnston et al., 2019). The Klf5 deletion in mouse lead intestine epithelial damage and a reduction of colon proliferative crypt cells (Nandan et al., 2014). Our analysis showed that TP63, TFAP2A, TFAP2C, and GRHL1 are common important TFs in the skin and esophagus (Figure 6B). The function of these TFs has been well studied in the skin. TP63 is one of the TFs that is important in both skin and esophagus development (Daniely et al., 2004; Kurita et al., 2018; Qu et al., 2018). TP63 and TFAP2A have been used in *in vivo* reprogramming of wound-resident cells to generate skin epithelial tissue (Kurita et al., 2018). Both TFAP2A and TFAP2C are required for proper early morphogenesis and development as well as terminal differentiation of the skin epidermis (Budirahardja et al., 2016; Kousa et al., 2018; Wang et al., 2008). GRHL1 is important for the functioning of the epidermis. Grhl1 knockout mice exhibit palmoplantar keratoderma, impaired hair anchoring, and desmosomal abnormalities (Wilanowski et al., 2008). It would be interesting to investigate what roles they play in esophagus. PAX9 regulates squamous cell differentiation and carcinogenesis in the oro-oesophageal epithelium (Xiong et al., 2018). Although not all predicted TFs are known to have an important role in specific tissues, further research is warranted. The TFs in the TF atlas predicted by ANANSE may also be good candidates for studying tissue development and engineering in regenerative medicine.

Another large benefit of the model that we implemented in ANANSE is the wide applicability. The source code of ANANSE is publicly available under a liberal license. ANANSE does not depend on large collections of reference data and it is straightforward to run on new data, such as different cell types or even species. The types of data required for this analysis are the following: gene expression data (RNA-seq) and genome-wide assays of enhancer activity. The enhancer data can be ATAC-seq, H3K27ac ChIP-seq or a combination of them, which is relatively easily obtained, not only in human cell types or in common model species, but also often in non-model species (Villar et al., 2015). The predictions of ANANSE, represented by TF binding, gene regulatory networks and TFs ranked by their influence score, are useful to study gene regulatory principles in a wide variety of contexts. While we used trans-differentiation experiments to benchmark the TF influence score, the utility of ANANSE is not limited to these types of experiments. It can also be used to study differentiation, cell type-specific gene regulation and developmental processes.

We also acknowledge limitations in our approach. In ANANSE, we link enhancer regions to genes on the basis of distance. For each TF and gene interaction pair, ANANSE only considers TF binding information located at most 100kb up and downstream of the corresponding gene. Although data from a recent CRISPR enhancer interference screen showed that genomic distance is largely informative in predicting enhancer-target interactions (Fulco et al., 2019), this approach may be limited when applying to genes regulated through ultra-long range regulation or through less abundant inter-chromosomal contacts (Olivares-Chauvet et al., 2016). This limitation of our method can potentially be addressed using chromosome conformation capture techniques (3C) (Dekker et al., 2002) or other adaptations as circular 3C (4C) (Simonis et al., 2006; Zhao et al., 2006), chromosome conformation capture carbon copy (5C) (Dostie et al., 2006), chromatin immunoprecipitation using PET (ChIA-PET) (Fullwood et al., 2009) and Hi-C (Lieberman-Aiden et al., 2009). However, these types of data are currently only available for a limited number of cell types, therefore incorporation of topology data would limit the broad utility and application of our approach. Another limitation is that the current implementation of ANANSE focuses on activating transcription factors. During cell differentiation and reprogramming, other factors such as transcriptional repressors and chromatin modifying enzymes also play an important role. These are currently not considered in ANANSE. Finally, similar to other genome-wide gene regulatory network inference methods, the performance of ANANSE is not yet optimal. While our benchmarks (Figure 4 A and B, Supplementary Figure S2) indicate that ANANSE improves upon other methods, it is clear that there is still much progress to be made in genome-wide GRN inference.

## Conclusion

Here we presented ANANSE, a computational tool for 1) transcription factor binding prediction, 2) gene regulatory network inference and 3) efficient prediction of TFs in cell fate determination. It outperforms other published methods in GRN inference and in predicting TFs that can induce trans-differentiation. In addition, it is open source, freely available and can be easily applied to other cell types and in any species. In summary, ANANSE exploits the powerful impact enhancers have on gene regulatory networks, and it provides insights into TF mediated regulatory mechanisms underlying cell fate determination and development.

## Supplementary information

**Additional file 1: Figure S1.** Comparison of ANANSE TF binding prediction performance to other methods. **Figure S2.** Evaluation of tissue-specific enhancer GRNs predicted by ANANSE. **Figure S3.** Evaluation of predicting tissue-specific GRNs based on enhancer, promoter or expression data. **Figure S4.** Evaluation of tissue-specific enhancer GRNs with enhancer-gene summarization methods. **Figure S5.** Comparison of the top 10 key TFs predicted by different GRN sizes in seven experimentally validated trans-differentiation strategies. **Figure S6.** Comparison of different GRN sizes used in the ANANSE prediction in seven experimentally validated trans-differentiation strategies. **Figure S7.** Comparison of the top 10 key TFs predicted by different GRNs (enhancer, promoter and expression) in seven experimentally validated trans-differentiation strategies. **Figure S8.** Evaluation of the performance of ANANSE using experimentally validated trans-differentiation strategies. **Figure S9.** Comparison of the top 10 key TFs predicted by different methods in seven experimentally validated trans-differentiation strategies. **Figure S10.** Classification of the human gene expression and TF influence score.

**Additional file 2: Table S1.** Overview of accessions of public data that was used.

**Additional file 3: Table S2.** Summary of genomic locations of TF binding sites

**Additional file 4: Table S3.** Trans-differentiation expression dataset

**Additional file 5: Table S4.** Differential gene expression results for cell types used the trans-differentiation benchmark.

**Additional file 6: Table S5.** Trans-differentiation key TFs prediction results among different network sizes

**Additional file 7: Table S6.** Trans-differentiation key TFs prediction results among enhancer promoter and expression network

**Additional file 8: Table S7.** Application of ANANSE to human tissue data

**Additional file 9: Supplementary Note S1**. Detailed description of the GRN benchmarking procedure.

## Acknowledgements

This study used data provided by the ENCODE consortium (https://www.encodeproject.org/) and the NIH Roadmap Epigenomics Consortium (http://nihroadmap.nih.gov/epigenomics/). We would like to thank Jos Smits, Branco Heuts, Jori de Leuw and Seline van den Oever for providing critical input during the development of ANANSE.

## Funding

QX was funded by the Chinese Scholarship Council (grant 201606230213). SJvH was supported by the Netherlands Organization for Scientific Research (NWO grant 016.Vidi.189.081). Early work on this project was supported by a US National Institutes of Health grant (NICHD, R01HD069344) to GJCV.

*Conflict of interest statement.* None declared.

---

**Supplementary Figure S1.**
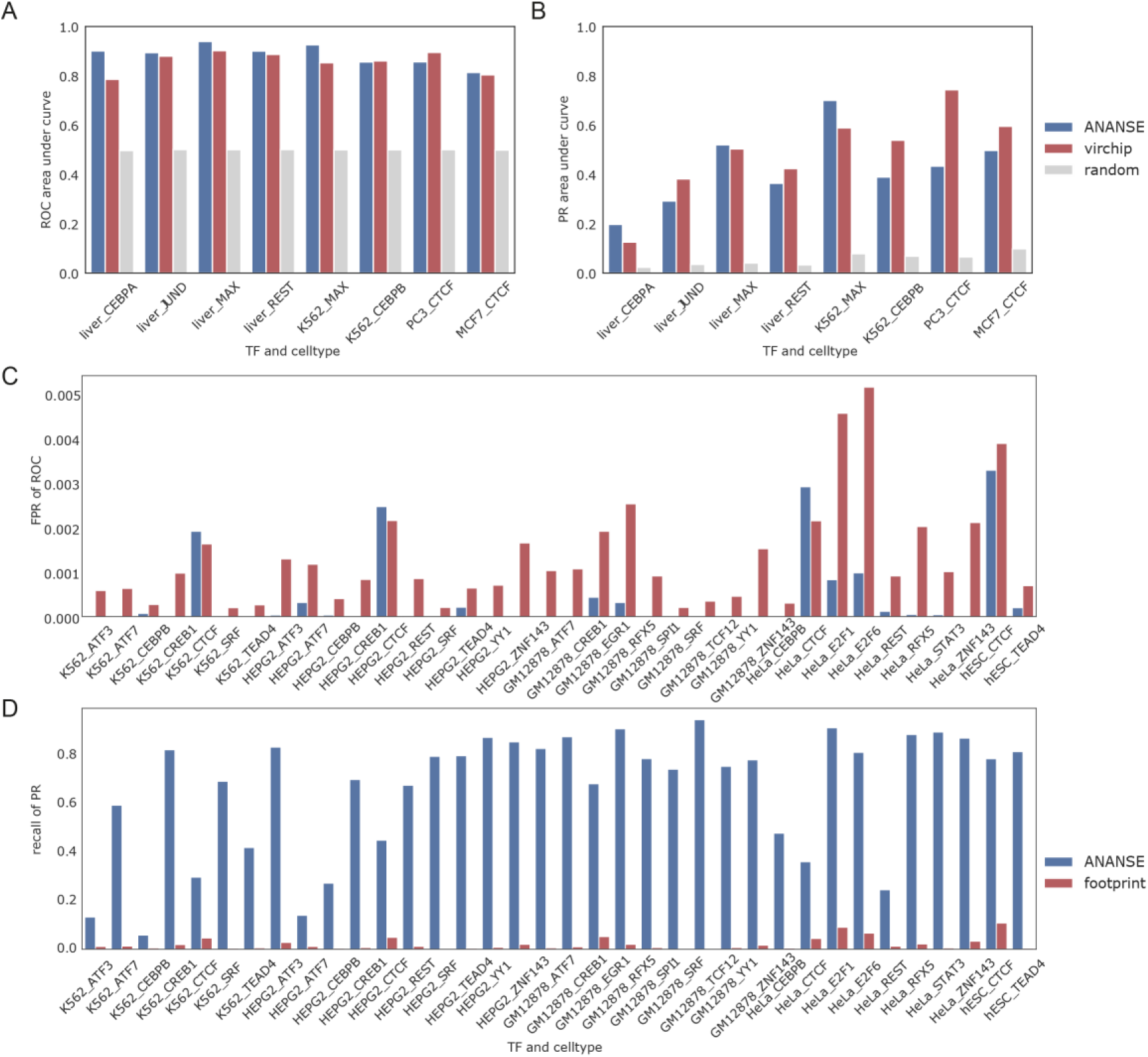
Comparison of ANANSE TF binding prediction performance to other methods. **(A)** The barplot shows ROC AUC of TF binding prediction of ANANSE compared to the Virtual ChIP-seq predictions in the ENCODE-DREAM validation cell types. ROC AUC of the ANANSE predicted TF binding is shown in blue; the Virtual ChIP-seq predicted result in red and the random model is indicated in gray. **(B)** The same evaluation as in A), with the PR AUC shown as a barplot. **(C)** Comparison of ANANSE binding predictions to DNAseI footprints obtained from (Vierstra et al., 2020). The barplot shows False Positive Rate (FPR) of TF binding prediction of ANANSE (blue) compared to the footprint TF binding prediction (ref) in ENCODE-DREAM validation cell types. The FPR is shown at the selected TPR of the footprints. **(D)** The same evaluation as in C), with the Recall shown as a barplot. The recall for ANANSE is calculated at the same precision of the footprint-based TF binding prediction.

**Supplementary Figure S2.**
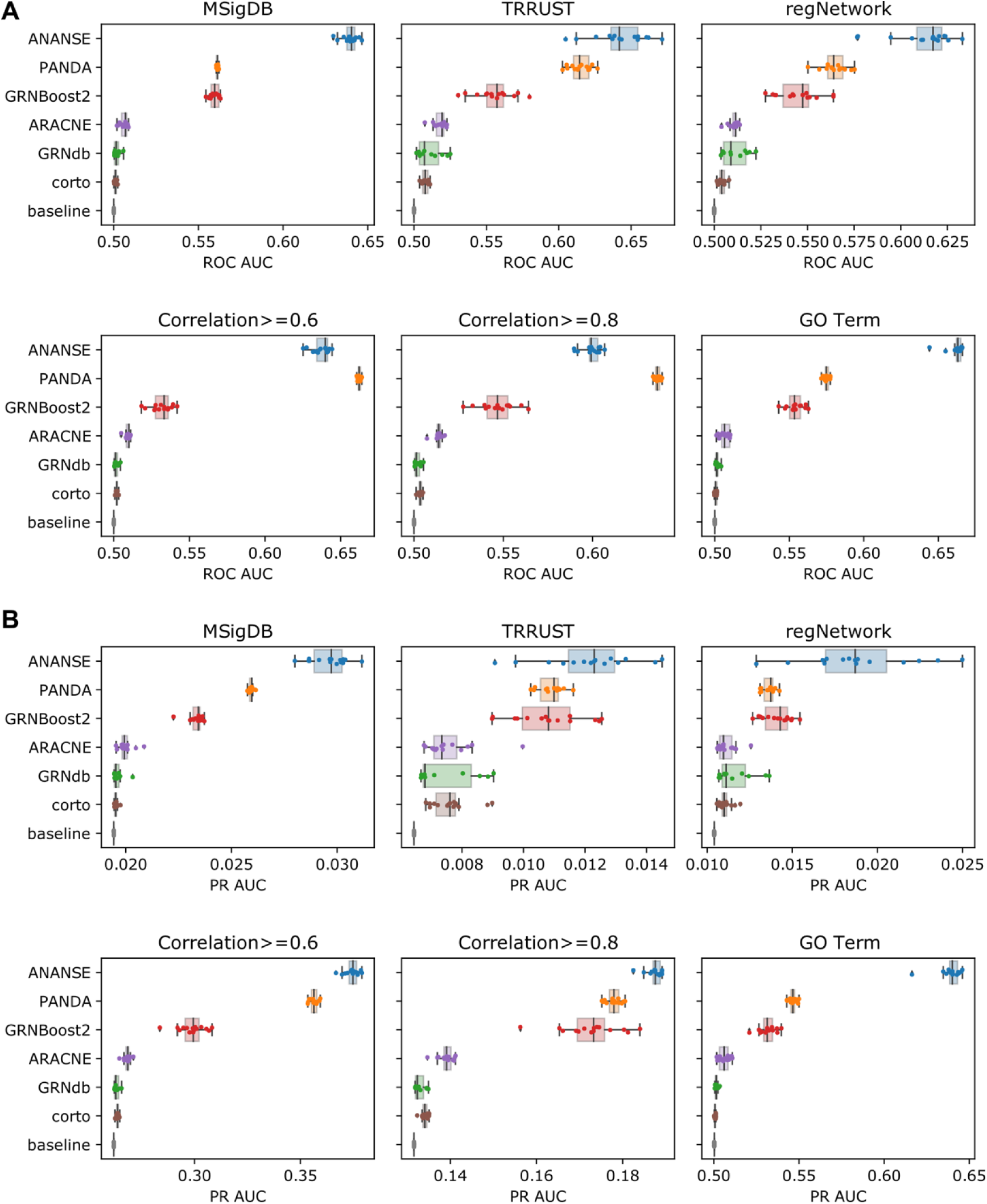
Evaluation of tissue-specific enhancer GRNs predicted by ANANSE. (A) Evaluation of the predicted networks, similar to Figure 4A and 4B, using different types of data: two TF-Gene regulatory networks based on interaction databases (MsigDB, TRRUST, regNetwork, Correlation>=0.6, Correlation>=0.8, and GO Term). The boxplots show the AUC of ROC for 15 different tissues. ROC AUC of the ANANSE predicted networks is shown in blue; the PANDA networks in orange; the GRNdb networks in green; the GRNBoost2 networks in red; the ARACNE networks in puple; the corto networks in brown; the random baseline networks in gray. (B) The same evaluation as in A), with the PR AUC shown as a boxplot.

**Supplementary Figure S3.**
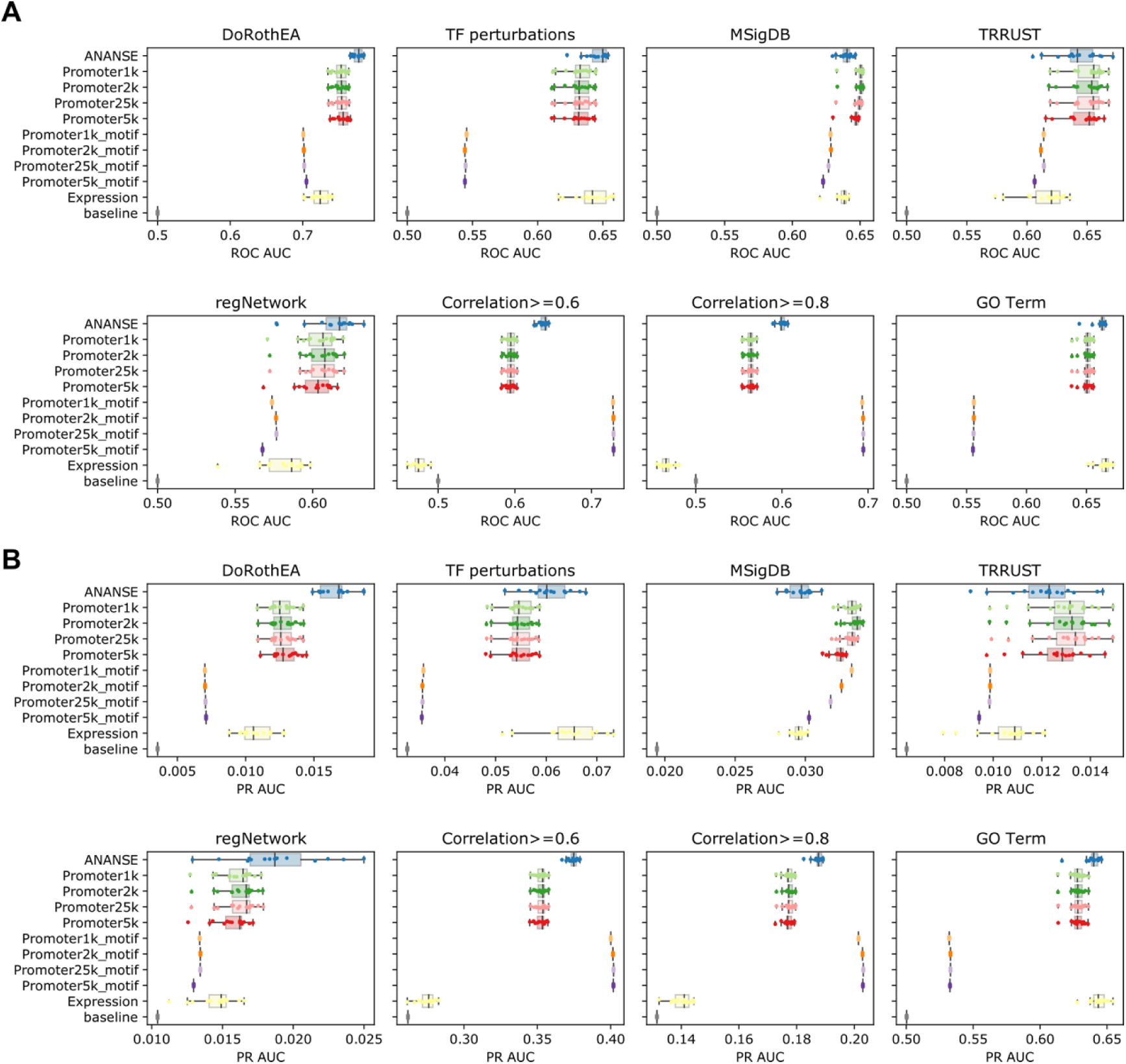
Evaluation of predicting tissue-specific GRNs based on enhancer, promoter or expression data. (A) The boxplots show the evaluation of four types of networks: ANANSE (the full model), PromoterXK (ANANSE based on promoters, instead of enhancers), PromoterXK_motif (GRN inference based on motif score alone) and Expression (GRN inference based on expression of TF and target gene alone). Four definition of promoters were tested, based on the distance around the gene TSS: 1kb, 2kb, 5kb and 25kb. The networks were evaluated based on the same references as Fig. 4 and Sup. Fig S2. Shown is the ROC AUC of the tissue-specific networks compared to the reference data. (B) The same evaluation as in A), with the PR AUC shown as a boxplot.

**Supplementary Figure S4.**
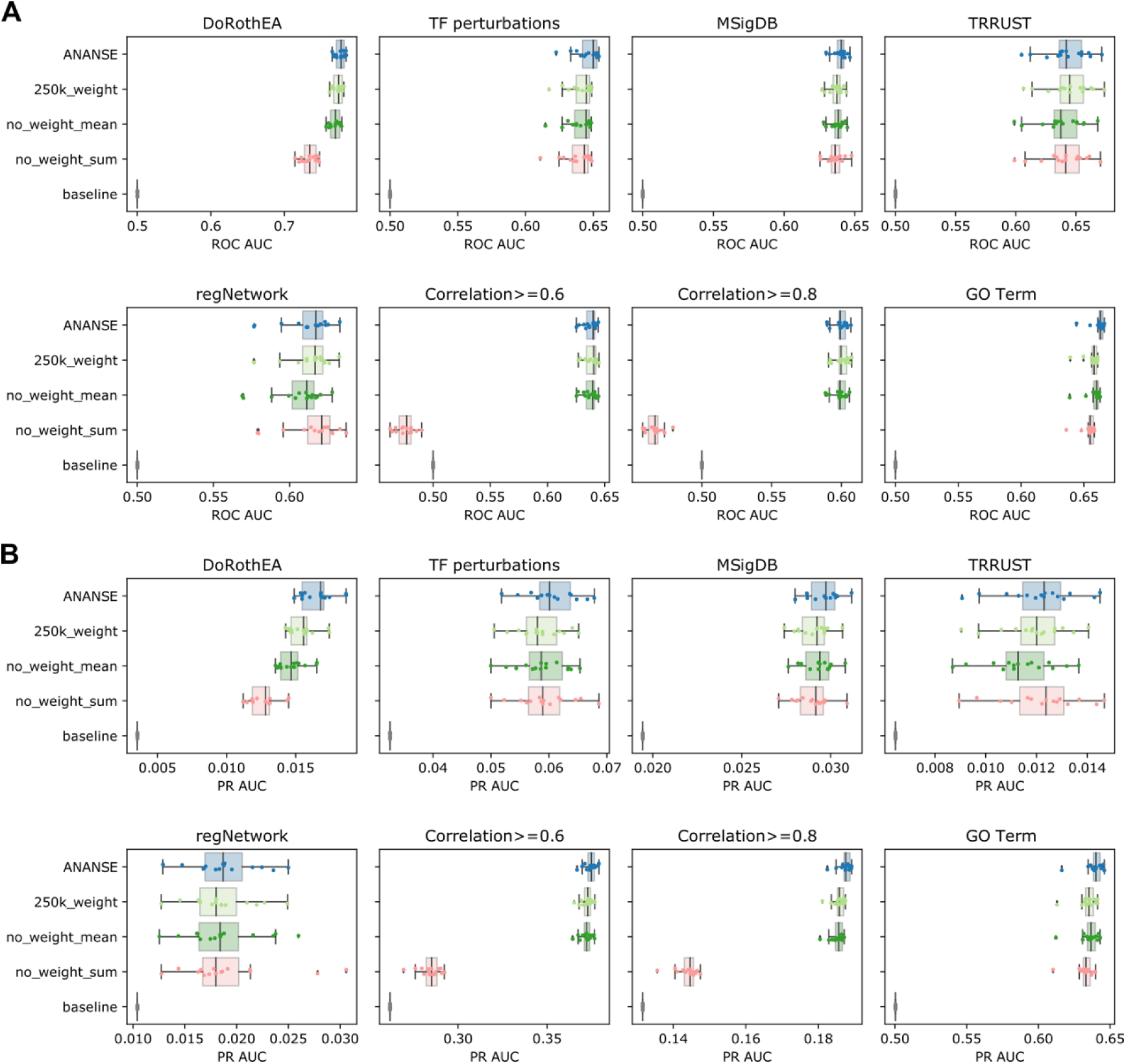
Evaluation of tissue-specific enhancer GRNs with enhancer-gene summarization methods. (A) Evaluation of the predicted GRNs, as described before. The plots show four different methods of calculating the TF-gene binding score: ANANSE, 250k_weight (the distance-weighted sum with enhancers within 250kb instead of 100kb of the gene TSS), no_weight_mean (the mean of all scores within 100kb, with equal weight) and no_weight_sum (the sum of all scores within 100kb, with equal weight). The boxplots show the AUC of ROC for 15 different tissues. ROC AUC of the ANANSE predicted networks is shown in blue; the 250kb weighted enhancer networks in light green; the 100kb mean enhancer networks in green; the 100kb sum enhancer networks in light red; the random baseline networks in gray. (B) The same evaluation as in A), with the PR AUC shown as a boxplot.

**Supplementary Figure S5.**
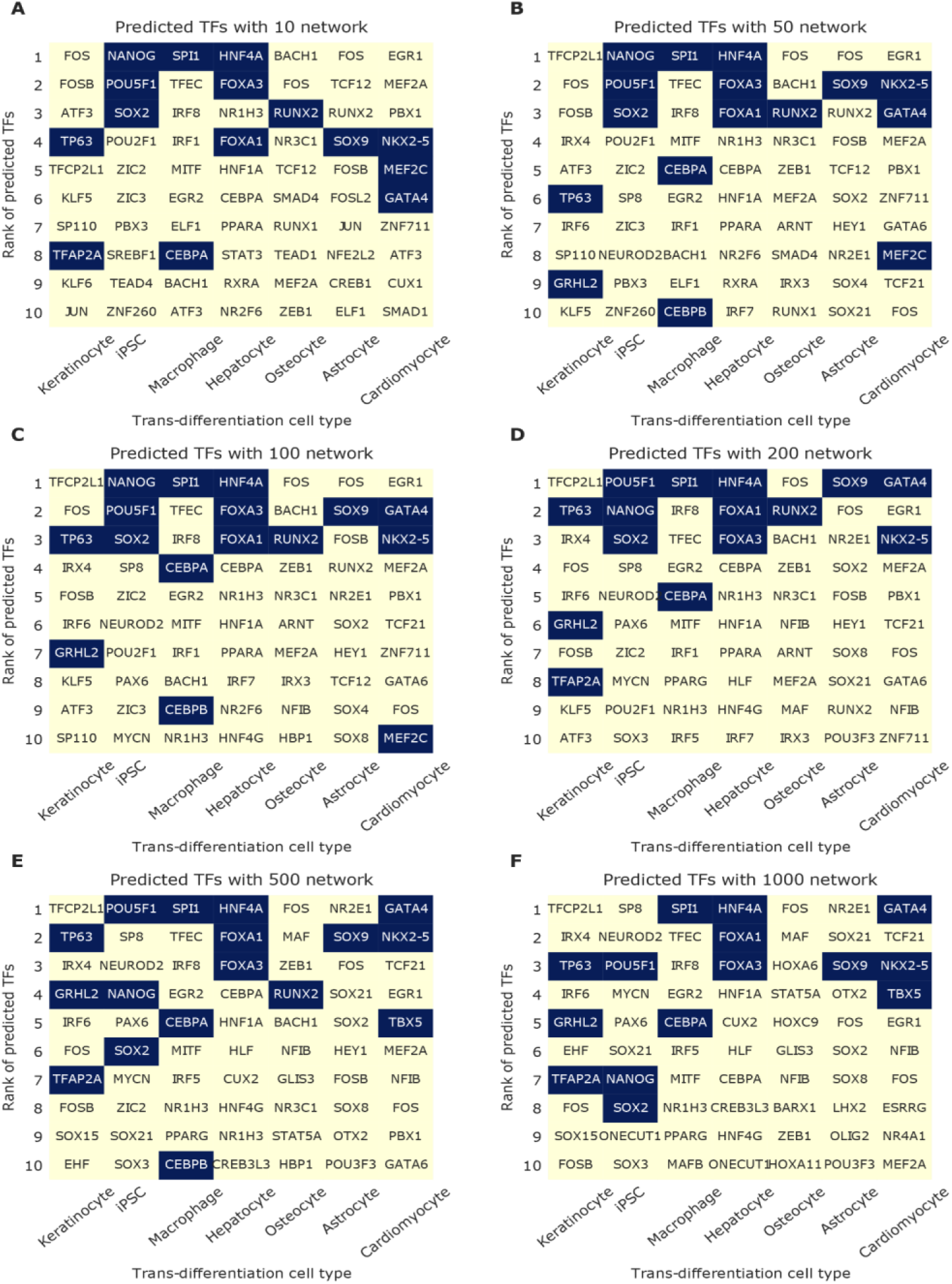
Comparison of the top 10 key TFs predicted by different GRN sizes in seven experimentally validated trans-differentiation strategies. The x-axis shows seven experimentally validated trans-differentiations, and the y-axis shows the top 10 predicted key TFs ranked by their influence score. Black boxes highlight the TFs that were used in trans-differentiation experiments. **(A)** 10k network. **(B)** 50k network. **(C)** 100k network. **(D)** 200k network. **(E)** 500k network. **(F)** 1000k network.

**Supplementary Figure S6.**
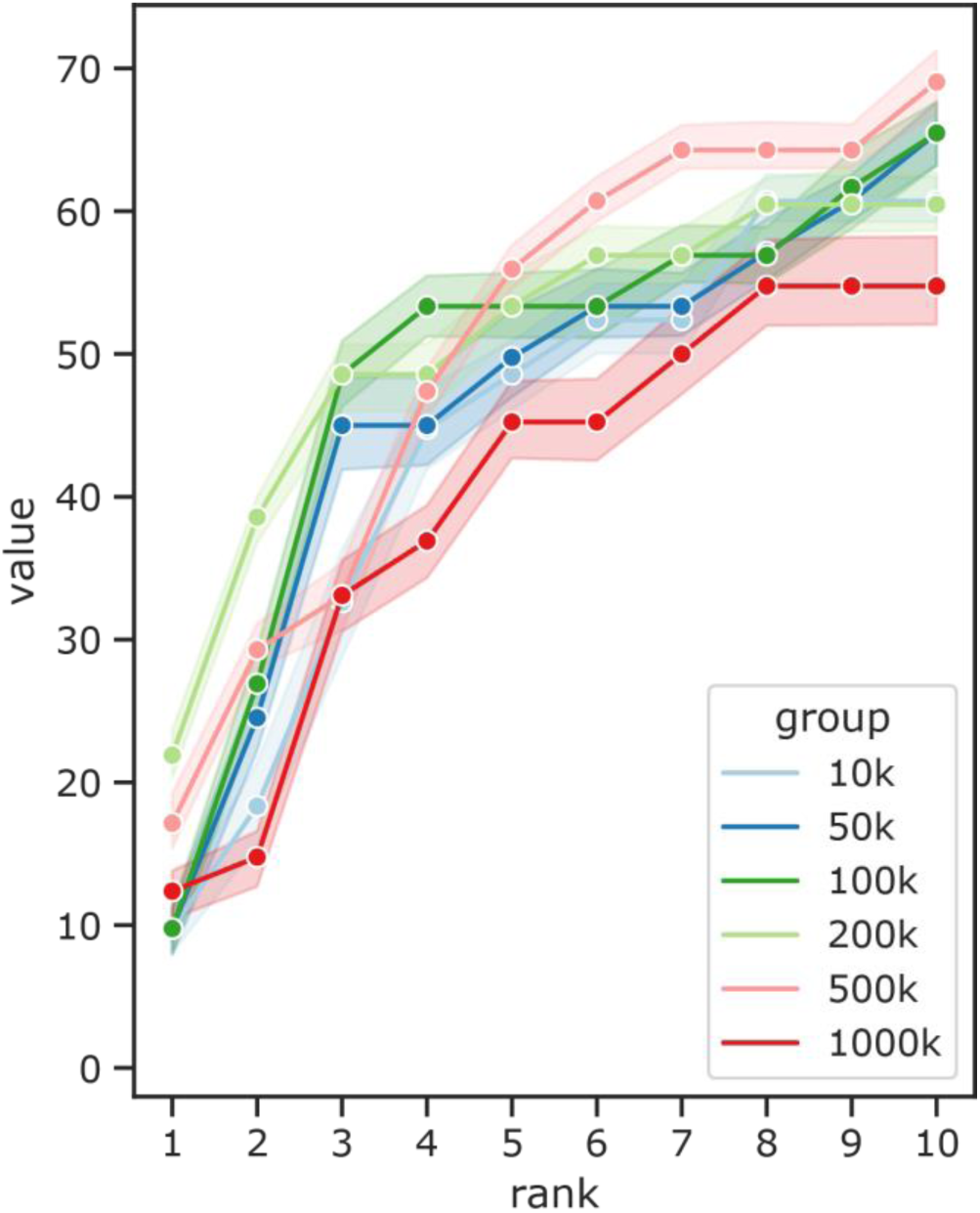
Comparison of different GRN sizes used in the ANANSE prediction in seven experimentally validated trans-differentiation strategies. **(A)** The line plots show the comparison of the predicted key TFs for six different sizes of GRNs Shown is the fraction of predicted TFs compared to all known TFs based on trans-differentiation protocols described in the literature (y-axis) as a function of the top number of TFs selected (x-axis). The shaded area represents the minimum and maximum percentage of corresponding recovered TFs when using six out of seven trans-differentiations.

**Supplementary Figure S7.**
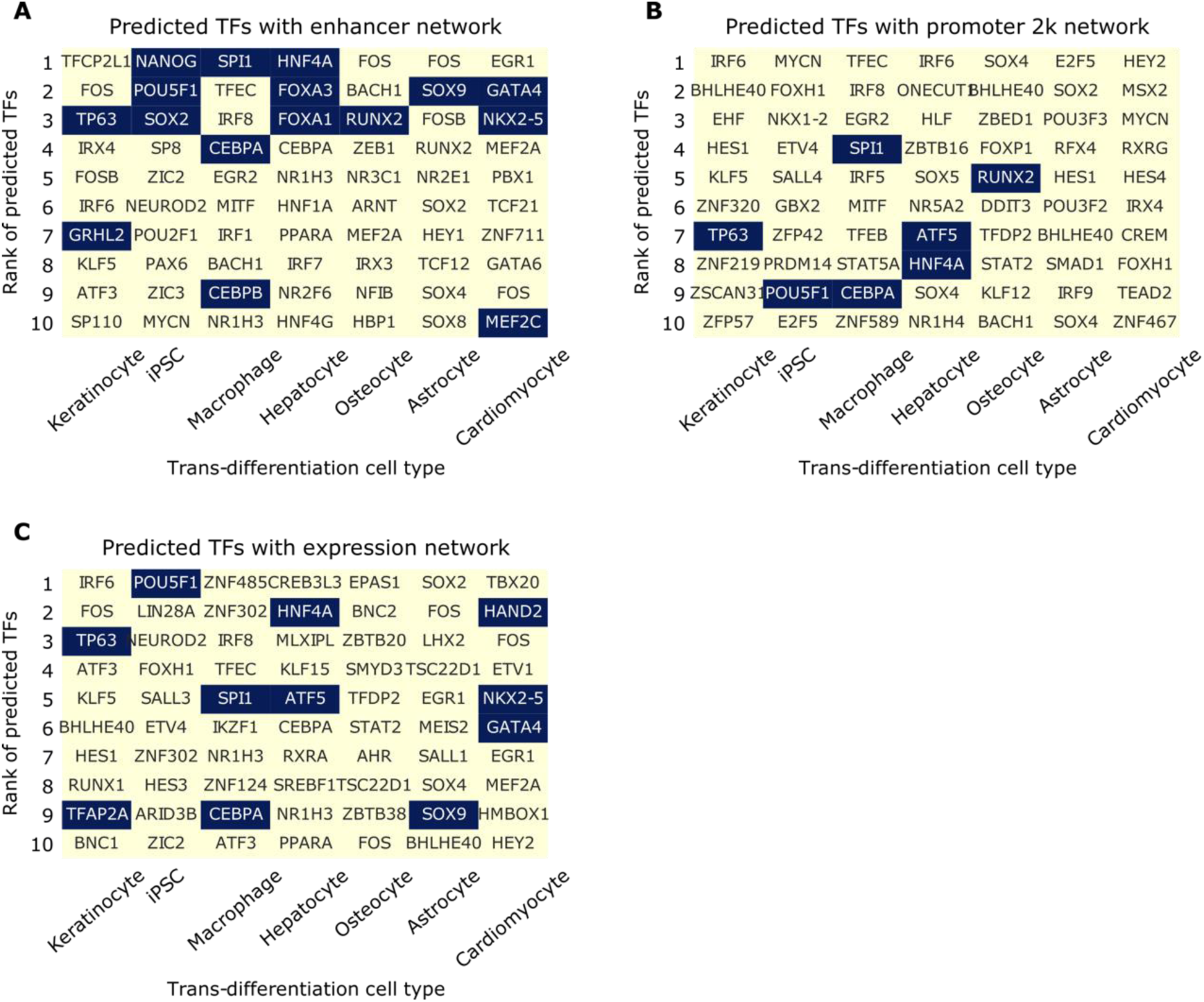
Comparison of the top 10 key TFs predicted by different GRNs (enhancer, promoter and expression) in seven experimentally validated trans-differentiation strategies. The x-axis shows seven experimentally validated trans-differentiations, and the y-axis shows the top 10 predicted key TFs ranked by their influence score. Black boxes highlight the TFs that were used in trans-differentiation experiments. **(A)** The results for ANANSE, based on a GRN that was inferred using peaks, regardless of their genomic location (includes both promoters and enhancers). **(B)** The results for ANANSE, based on a GRN that was inferred using the highest peak in the gene promoter, defined as < 2kb from the gene transcription start site. **(C)** The results for ANANSE, based on a GRN that was inferred using only the expression levels of the TFs and target genes.

**Supplementary Figure S8.**
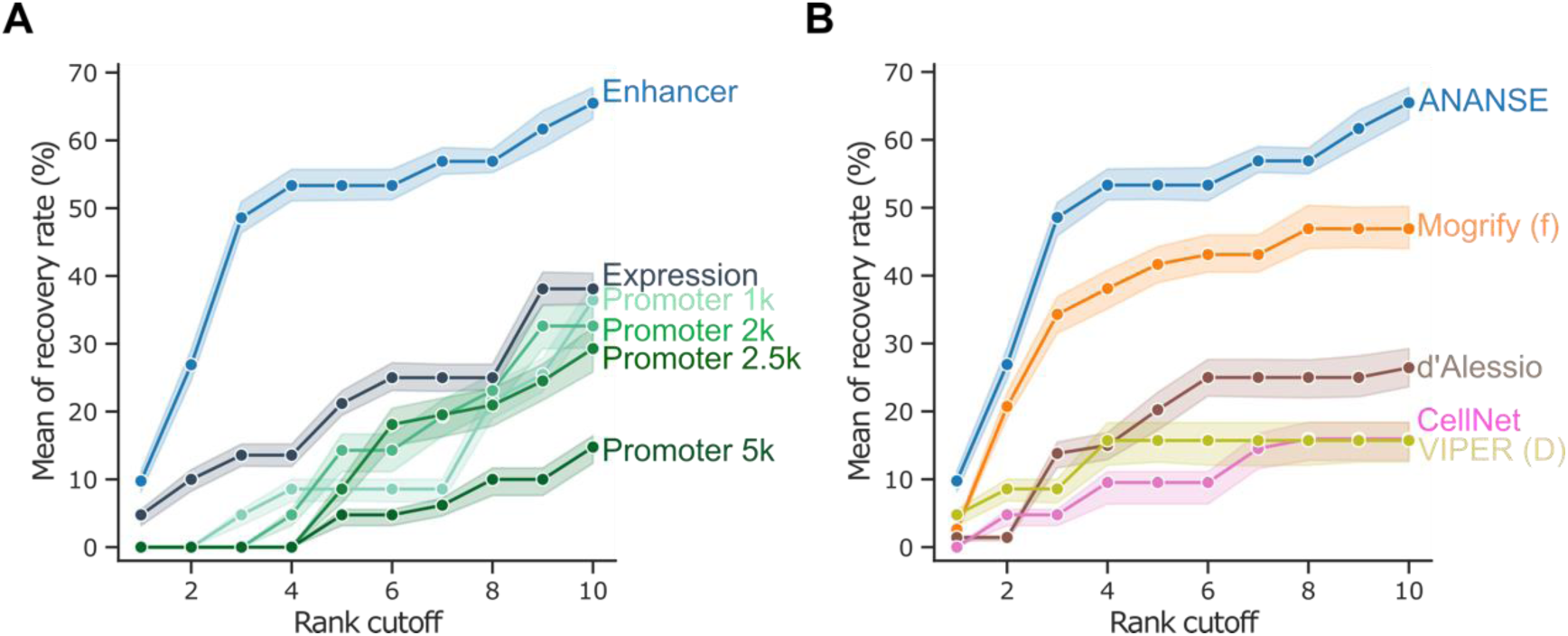
Evaluation of the performance of ANANSE using experimentally validated trans-differentiation strategies. (A) The line plots show the comparison of the predicted top TFs for trans-differentiation from cell type-specific networks. Based on the difference between two networks, TFs were prioritized using the influence score calculation implemented in ANANSE. Shown is the fraction of predicted TFs compared to all known TFs based on trans-differentiation protocols described in the literature (y-axis) as a function of the top number of TFs selected (x-axis). The mean of recovery rate is the average of all TF sets when corresponding trans-differentiation has server different experimental validate TF sets. The shaded area represents the minimum and maximum percentage of corresponding recovered TFs when using six out of seven trans-differentiations. Three different types of networks were used: gene expression (deep green), promoter-based (promoter in 1kb, 2kb, 2.5kb, and 5kb) TF binding in combination with expression (blue), and enhancer-based TF binding in combination with expression (blue). (B) The line plots show the comparison of the predicted top TFs for trans-differentiation based on different computational methods. The y-axis indicates the percentage of experimentally validated cell TFs that are recovered as a function of the number of top predictions, similar as in A). Six different methods are shown: ANANSE (blue), Mogrify (full) (orange), d’Alessio (brown), CellNet (red), and VIPER with Dorothea network (yellow). The shaded area represents the minimum and maximum percentage of corresponding recovered TFs when using six out of seven trans-differentiations. CellNet only contains data from fibroblast to ESC, Hepatocyte, and Macrophage; and Mogrify and CellNet only contain the top 8 predicted factors.

**Supplementary Figure S9.**
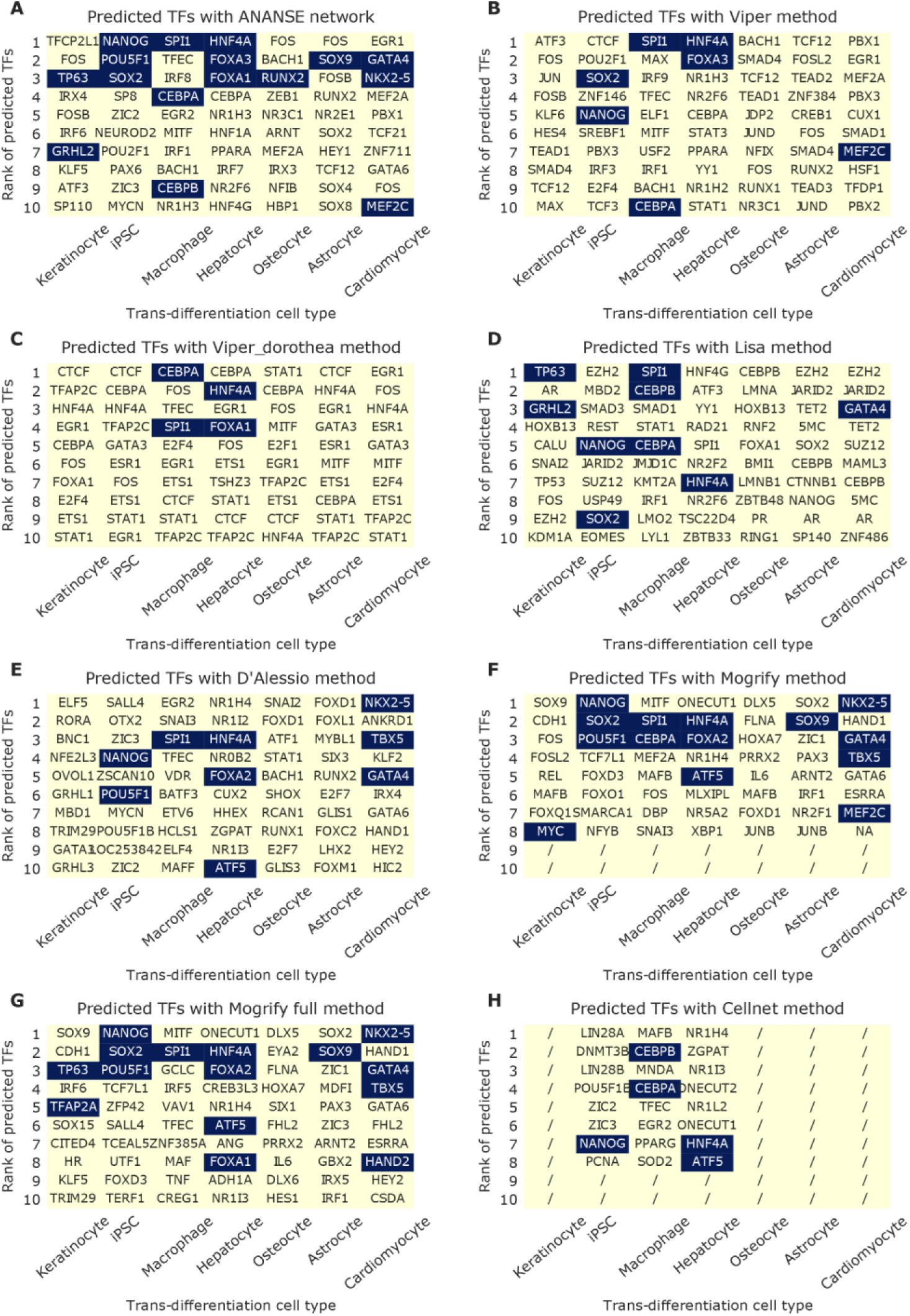
Comparison of the top 10 key TFs predicted by different methods in seven experimentally validated trans-differentiation strategies. The x-axis shows seven experimentally validated trans-differentiations, and the y-axis shows the top 10 predicted key TFs ranked by their influence score. Black boxes highlight the TFs that were used in trans-differentiation experiments. **(A)** ANANSE. **(B)** Lisa. **(C)** D’Alessio. **(D)** Mogrify. **(E)** Mogrify full list. **(F)** CellNet.

**Supplementary Figure S10.**
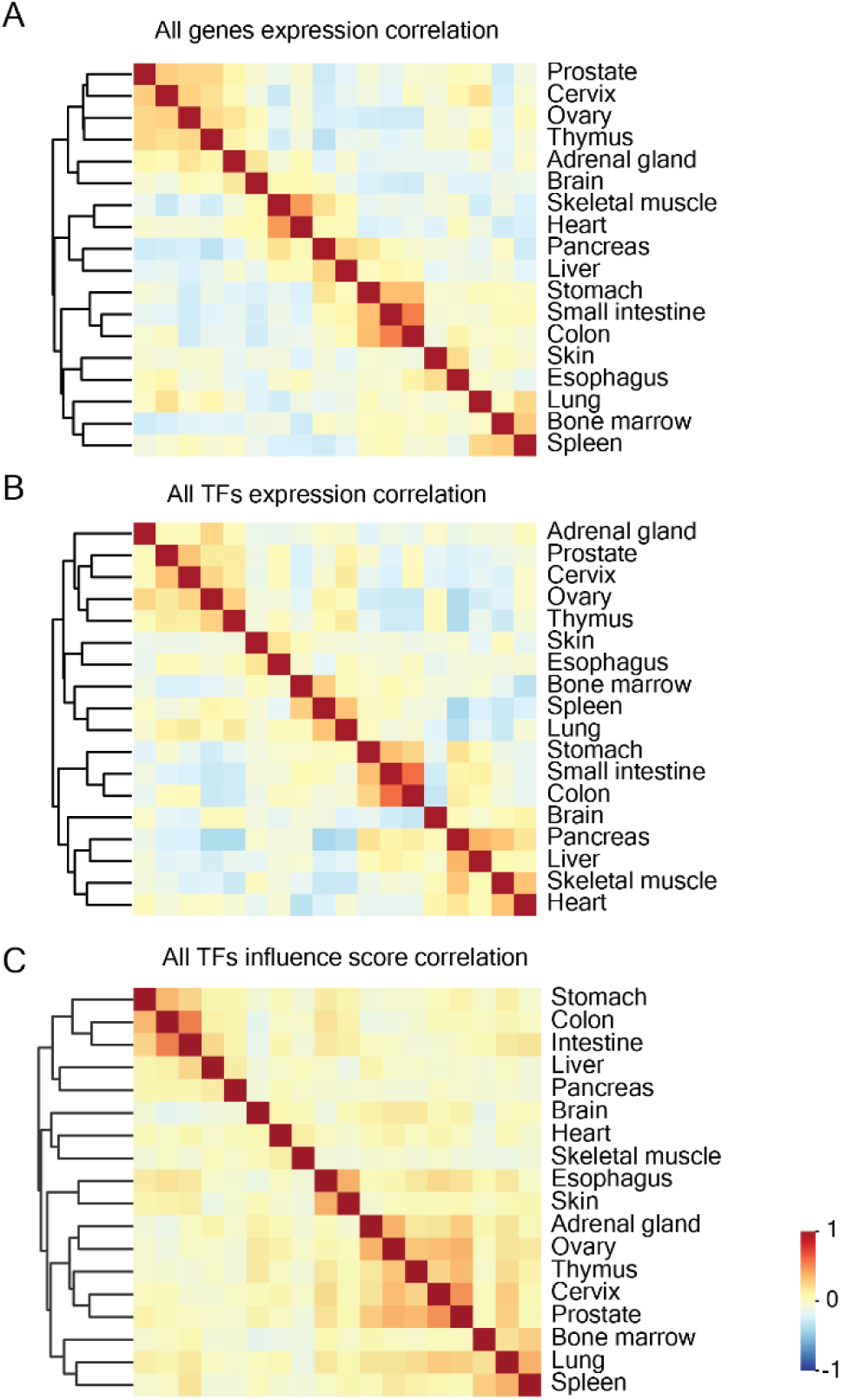
Classification of the human gene expression and TF influence score. **(A)**, Heatmap showing the pairwise correlation between all 18 tissues based on gene expression. The colors in the heatmap indicate high (red) or low (blue) correlation across the tissue set. **(B)**, Heatmap showing the pairwise correlation between all 18 tissues based on the expression of all TFs. **(C)**, Heatmap showing the pairwise correlation between all 18 tissues based on TF influence scores.

## Notes

https://doi.org/10.5281/zenodo.4066423

https://doi.org/10.5281/zenodo.4814015

https://doi.org/10.5281/zenodo.4809062

https://doi.org/10.5281/zenodo.4814016

https://github.com/vanheeringen-lab/ANANSE

https://github.com/vanheeringen-lab/ANANSE-manuscript

### Competing Interest Statement

The authors have declared no competing interest.

### Summary of Updates

First, we have improved the prediction of transcription factor binding by using a more detailed model: 1) We now use ATAC-seq as well as H3K27ac ChIP-seq data for the model. This results in improved performance (see Figure 3A in the revised manuscript). Although the combined data works best, a model based on either ATAC-seq alone or H3K27ac ChIP-seq alone can also be used. This means that the approach is more widely applicable, even when only one of the two data types is available. 2) We have trained transcription factor specific models based on all available TF ChIP-seq data (237 TFs) in the REMAP project for 6 commonly used cell lines. A non-specific model (trained on all TFs, comparable to the model in the previous ANANSE version) is available as fallback, for when ChIP-seq of a TF was not available for training. The estimation of the binding performance is now based on these 237 TFs in the 6 cell lines. In addition, the ROC AUC and PR AUC metrics are performed on held-out chromosomes, in held-out cell types. Second, we have added inferred motif activity to the network inference. This, in combination with the improved binding prediction, results in a greatly improved GRN inference performance. We have revisited all benchmarks. We added DoRothEA and TF perturbation references, and now include two additional state-of-the-art reference methods: GRNBoost2 (a GENIE3-like approach) and networks from GRNdb that were inferred on single-cell RNA-seq using SCENIC (Figure 4A and B).

