## Supplementary Note S1 for "ANANSE: An enhancer network-based computational approach for predicting key transcription factors in cell fate determination"

**Supplemental Note S1: GRN benchmarking**

One of the options to benchmark inferred gene regulatory networks (GRNs) is to compare the inferred interaction scores (edge strength) to a reference database (or GRN) containing known regulatory interactions. There are a few caveats to this approach. First, most reference databases contain a limited number of interactions. This means that are a relatively large number of false negatives, i.e. TF-target interactions that are not in the database. Second, the reference database and the inferred GRN may contain TFs or target genes that are present in one, but not in the other. For instance, the reference database will likely not contain interactions for all transcription factors. Other issues may occur, for instance, due to incompatible and/or incomplete gene symbols. To benchmark GRNs in this manuscript, we have used the following approach.

As reference, we take the Cartesian product, $T_{ref}\times G_{ref}$, where $T_{ref}$is the set of all transcription factors in the reference and $G_{ref}$ is the set of all target genes in the reference (which may also include transcription factors). For all combinations of $t \in T_{ref}$ and $g \in G_{ref}$ we set the interaction score $i_{t,g}$ to 1 if it is present in the reference, and 0 otherwise. We then compare all these interactions to the edges in the GRN that we want to benchmark. If a specific interaction $i_{t,g}$is not present in that GRN, we set the score to the minimum score of the GRN.
Here is a more visual example.

Database containing regulatory interactions:

```
TF	Target
TF1	GENE1
TF2	GENE2
TF3	GENE1
TF4	GENE3
```

Constructed reference network:

```
TF	Target	Interaction
TF1	GENE1	1
TF1	GENE2	0
TF1	GENE3	0
TF2	GENE1	0
TF2	GENE2	1
TF2	GENE3	0
TF3	GENE1	1
TF3	GENE2	0
TF3	GENE3	0
TF4	GENE1	0
TF4	GENE2	0
TF4	GENE3	1
```

Let’s say we compare against the following inferred GRN (note that TF3 and TF4 are missing in this analysis):

```
TF	Target	Interaction
TF1	GENE1	0.9
TF1	GENE2	0.2
TF1	GENE3	0.1
TF2	GENE1	0.5
TF2	GENE2	0.8
TF2	GENE3	0.3
```

We now join the inferred GRN to the reference network (which would be a left join in pandas/SQL terms):

```
TF	Target	Ref_int	GRN_int
TF1	GENE1	1		0.9
TF1	GENE2	0		0.2
TF1	GENE3	0		0.1
TF2	GENE1	0		0.5
TF2	GENE2	1		0.8
TF2	GENE3	0		0.3
TF3	GENE1	1		NA
TF3	GENE2	0		NA
TF3	GENE3	0		NA
TF4	GENE1	0		NA
TF4	GENE2	0		NA
TF4	GENE3	1		NA
```

All missing interaction score are now set to a value lower than the lowest score in the network:

```
TF	Target	Ref_int	GRN_int
TF1	GENE1	1		0.9
TF1	GENE2	0		0.2
TF1	GENE3	0		0.1
TF2	GENE1	0		0.5
TF2	GENE2	1		0.8
TF2	GENE3	0		0.3
TF3	GENE1	1		0.0
TF3	GENE2	0		0.0
TF3	GENE3	0		0.0
TF4	GENE1	0		0.0
TF4	GENE2	0		0.0
TF4	GENE3	1		0.0
```

Using Ref_int as y_true and GRN_int as y_score, we can now calculate metrics such as the area under the curve of the ROC curve or the area under the curve of the Precision-Recall curve. Note that $T_{ref}\times G_{ref}$ is large (~1500 TFs x ~20,000 genes), while the number of regulatory edges in reference databases is relatively small. This means that the dataset is highly unbalanced and the PR AUC is a better evaluation metric than the ROC AUC (Saito & Rehmsmeier, 2015).

An alternative to the approach described here is to take the intersection of the reference and the inferred GRN, $(T_{ref}\times G_{ref}) \cap{(T}_{GRN}\times G_{GRN})$. However, this does not result in a fair comparison between GRNs, as the number of TFs and target genes that is reported as output of a GRN inference method may be very different.
